# Human parainfluenza virus infection remodels the host cell glycome

**DOI:** 10.1101/2025.06.15.659808

**Authors:** Plabon Kumar Das, Larissa Dirr, Benjamin Bailly, Patrice Guillon, Arun Everest-Dass, Mark von Itzstein

## Abstract

Human parainfluenza virus type 3 (HPIV-3) remains a major cause of respiratory illness particularly among young children, the elderly and immunocompromised individuals. Despite significant efforts in therapeutic discovery research, there is neither an effective antiviral nor a vaccine available against HPIV-3. Host cell glycosylation is known to play a pivotal role in virus entry and replication. While some host glycan-based cellular receptors for HPIV-3 have been identified, the dynamics of the host glycome upon HPIV-3 infection has never been studied. Herein, we report the first mass spectrometry-based study that provides direct insight into the remodelling of the human lung adenocarcinoma cell (A549) glycome upon HPIV-3 infection. In this study we observed that HPIV-3 infection led to significant host-cell glycome changes in both oligomannose and sialylated complex-type *N-*glycans. Moreover, notable changes were also observed in both core 1 and core 2 type *O-*glycans, along with distinct glycosphingolipid remodelling in infected cells compared to their mock-infected counterparts. Our study presents the first evidence that hPIV-3 infection alters host-cell glycome, offering new insights into the virus’s impact on host cellular processes.

## Introduction

Human parainfluenza viruses (HPIVs) are among the most common causes of respiratory tract infections in children under the age of five, the elderly, and immunocompromised individuals, often resulting in high mortality rates^1^. The most common clinical manifestations reported for HPIV infections include colds, croup, bronchiolitis, bronchitis, and pneumonia. HPIV infections account for 40% of paediatric hospitalizations for lower respiratory tract infection and 75% of croup cases across the world, annually^1^. Despite the significant health burden posed by HPIV infections, neither antiviral drugs nor vaccines are currently available. The infection begins in the nose and oropharynx and then spreads to the lower respiratory tract in severe cases, with ciliated epithelial cells of both the upper and lower respiratory tracts being the primary targets of HPIVs^2,3^. There are four major serotypes of HPIV, designated as serotypes 1 through 4. HPIV-3 is the most prevalent HPIV subtype worldwide, including Australia, causing seasonal outbreaks in spring mostly after influenza outbreaks^4^. Regardless of serotypes, cell surface carbohydrates (glycans) play an important role in the life cycle of HPIVs^1^. Glycans, carbohydrate-based biopolymers linked by glycosidic bonds, are present in all living organisms. They play significant roles in biology, including maintaining the cell’s physical and structural integrity, contributing to the formation of extracellular matrix, mediating signal transduction, assisting in protein folding, and facilitating infection by pathogens^5^. In particular, glycans play numerous roles in viral pathogenesis, including as host cell receptors for adhesion of the virus particle (e.g. influenza virus)^6^, protection of the antigenic epitope of viral glycoproteins from the host immune response (e.g. human immunodeficiency virus)^7^, and subversion of the immune response by acting as an antibody decoy (e.g. Lassa virus)^8^. Like influenza viruses, HPIV-3 recognizes sialic acids on the host cell via its haemagglutinin-neuraminidase (HN) protein. The neuraminidase activity of the HN protein enables the virus to remove sialic acid residues from glycoproteins and glycolipids on the host cell surface, thereby aiding the release of new virus particles from the infected host-cell^9^. While the important role of glycans for entry and release of HPIV is known, the dynamics of the host-cell glycome expression upon infection remains to be determined. Recent advancements in chromatographic separation, ionization technologies, and mass spectrometry have substantially improved glycomic analyses, enabling high-throughput and structurally informative characterization of glycan species^10^. Notably, liquid chromatography (LC) employing porous graphitized carbon (PGC) stationary phases, coupled with electrospray ionization tandem mass spectrometry (PGC-LC-ESI-MS/MS), allows for precise separation and identification of glycan structural isomers based on retention time and fragmentation patterns¹¹. Using this PGC-LC-ESI-MS/MS workflow, we aimed to delineate the dynamic alterations in the host-cell glycome, including *N*-glycans, *O*-glycans, and glycosphingolipid (GSL) glycans, during HPIV-3 infection in human lung epithelial cells (A549).

Our results reveal extensive remodeling of the host-cell glycome following HPIV-3 infection at both 24 h and 48 h post-infection. Specifically, we observed a pronounced increase in oligomannose-type *N*-glycans and a concomitant reduction in sialylated complex-type *N*-glycans relative to mock-infected controls. Additionally, HPIV-3 infection induced higher expression of core 1 *O*-glycans alongside reduced levels of core 2 *O*-glycans. These changes in *N*- and *O*-glycan profiles were accompanied by significant alterations in GSL glycan composition. Collectively, our findings demonstrate that HPIV-3 orchestrates substantial glycome remodeling in host cells, providing the first comprehensive characterization of these glycomic shifts and offering new insights into HPIV-3 pathogenesis and potential therapeutic targets.

## Experimental procedures

### Cell culture, virus and infection assays

Human lung adenocarcinoma epithelial cells (A549; ATCC, CCL-185) were maintained in Dulbecco’s Modified Eagle Medium (DMEM) (Gibco, Cat. no. 10313021) media supplemented with 5% foetal bovine serum (FBS, Gibco, Cat. no. 10099141), 1% Antibiotic-Antimycotic (100X, Gibco, Cat. no. 15240062), 1% MEM Non-essential Amino Acid (100X, Sigma, Cat. no. M7145), 1% L-Glutamine 200 mM (100X, Gibco, Cat. no. 25030081) and incubated at 37 °C in a humidified chamber containing 5% CO_2_. Human parainfluenza virus type 3 (HPIV-3 JS), genetically modified to incorporate green fluorescent protein (GFP), was obtained from ViraTree (Product # P323). Virus propagation was performed in Rhesus monkey kidney epithelial (LLC-MK2; ATCC, CCL-7) cells using DMEM media supplemented with 1% Antibiotic-Antimycotic, 1% MEM Non-essential Amino Acid, 1% L-Glutamine (DMEM_inf_) at 35 °C in a humidified chamber containing 5% CO_2_. Approximately 2 million A549 cells were seeded in a 75 cm^2^ flask 48 h prior infection. Cells were washed 2 times with phosphate buffer saline (PBS) and 5 times with DMEM_inf_. Cell monolayer was infected with HPIV-3 for 1 h at 35 °C and 5% CO_2_, with multiplicity of infection (MOI) of 3 in triplicates. The mock-infected control was treated the same way as their infected counterpart except for adding viral inoculum. Both infected and mock-infected cells were incubated for 24 h and 48 h at 35 °C in a humidified chamber containing 5% CO_2_.

### Protein and glycosphingolipid (GSL) extraction

Cell culture media from both infected and mock-infected control were collected and mixed with 5 volumes of ice-cold (CH_3_)_2_CO (Sigma-Aldrich, Cat no. 650501) before storing at −20 °C overnight to precipitate secreted proteins. Infected and mock-infected cells were first washed four times with PBS to remove any residual media, including viral progeny. The cells were then detached with a scrapper and lysed in 1X cold RIPA lysis buffer (Thermo Fisher, Cat. no. 89900) supplemented with 1X protease inhibitor (Roche, cat. no. 11697498001). The mixture was incubated overnight at 4 °C, followed by sonication for 30 min in an ultrasound sonicator (VEVOR Ultrasonic Cleaner). Proteins and glycosphingolipids from cell lysates were extracted through standard chloroform, methanol, and water extraction method described elsewhere^11^. Briefly, 550 µL of chloroform (Sigma-Aldrich, Cat no. 1.02445) were added to each sample and sonicated for 15 min, followed by addition of 350 µL of methanol (Merck, Cat. no. 1.06035). The cell lysates were incubated at room temperature for 15 min with gentle shaking, followed by centrifugation at 3000 x g for 20 min. Upper phase containing glycosphingolipid were transferred to clean tubes. A mixture of 400 µL of chloroform/methanol (2:1) and 400 µL of methanol/water (1:1) was added to the pellets and vortexed vigorously. The mixture was sonicated again for 15 min followed by centrifugation at 3000 x g for 20 min. Upper phase was collected and added to the previous collected sample. An additional 400 µL methanol/water (1:1) mixture was added to the initial chloroform/methanol (2:1) phase and the separation steps were repeated. Proteins were pelleted, by adding 750 µL methanol to the lower phase, followed by centrifugation at 10000 x g for 20 min. The supernatant containing lipids were combined with previously collected upper aqueous phase and subsequently dried using a speed vac concentrator (Thermo Scientific; Savant SPD131DDA). The protein pellets were air-dried and then resuspended in 100 µL 8 M urea. The secreted protein samples were centrifuged at 5000 x g for 40 min to pellet the proteins. Supernatant was decanted and the protein pellets were resuspended in 100 µL 8 M urea. Bicinchoninic acid (Thermo-Fisher, Cat. no. 23227) assay was performed to quantify the proteins. Proteins were reduced and alkylated by adding dithiothreitol (25 mM final concentration) at 60 °C for 1 h and Iodoacetamide (50 mM final concentration) (Merck, Cat. no. 16125), for 30 min in dark at room temperature.

### Release of *N*- and *O*-glycans

The glycomics workflow, including the release, reduction, desalting, and PGC clean-up of *N-*, *O*-, and GSL glycans from protein and lipid samples, was performed according to a previously published protocol^12^, unless otherwise stated. Dot blotting of proteins on polyvinylidene fluoride (PVDF) membrane (0.45 µm pore size, Millipore, Cat. no.05317) was performed before the release of *N*-, and *O*-glycans. PVDF membrane was first activated with ethanol (Merck, Cat. no. 459828). Approximately 30 µg of proteins were spotted on the membrane. Protein spots were stained using direct Blue (Sigma-Aldrich, Cat. no. 212407), excised, and transferred to a flat bottom polypropylene 96-well plate (Merck, Cat. no. CLS3364), blocked with 1% (w/v) polyvinylpyrrolidone (Sigma-Aldrich, Cat no. PVP40) in 50% (v/v) methanol and washed 3 times with 100 μL ultrapure water (Sigma-Aldrich, Cat no. 1.15333). The *N*-glycans were then released using PNGase F (1000 units for each reaction) (New England Biolabs, Cat. no. P0709L) for 16 h at 37 °C. The released *N*-glycans were collected after agitation of the sealed plate on a shaker. The PVDF membrane wells were then washed with 2 × 20 µL of water and collated with previously collected *N*-glycans. Subsequently, 10 µL of 100 mM ammonium acetate (pH 5) was added to the collected *N*-glycans, followed by incubation at room temperature for 1 h. The samples were then dried under vacuum and kept in the freezer until further use.

To release the *O*-glycans from the same PVDF membrane spot, 2.5 µL of methanol was added to each spot to pre-wet them. About 50 µL 0.5 M sodium borohydride in 100 mM potassium hydroxide was added in each well and the plate was sealed with parafilm to prevent evaporation. Then the plate was incubated for 18 h at 50 °C. The reaction was neutralized with 5 µL glacial acetic acid.

### Purification and enzymatic release of GSL glycans

Purification of GSLs was performed according to a previously described method^13^. In brief, the dried aqueous and lipid-containing phases were resuspended in 100 μL of methanol for 30 min on a shaker, followed by the addition of 100 μL of water. Then, the sample was loaded onto a tC18 RP cartridge (Waters, Cat. no. WAT036820), that was preconditioned with 1 mL of chloroform/methanol (2:1; v/v), 1 mL of methanol and 2 mL of methanol/water (1:1; v/v). After the addition of the sample, the cartridge was washed with 2 mL of methanol/water and the elution was performed with 2 mL of methanol and 2 mL of chloroform and methanol (2:1; v/v). The combined eluate was dried under vacuum and stored at −20 °C until further use. The enzymatic release of the GSL head group was performed by following manufacturer’s instructions (New England Biolabs, Cat. no. P0773S). In brief, the dried GSLs were resuspended in 36 µL of WATER, 4 µL of EGCase I reaction buffer and 2 µL of EGCase I enzyme. The reaction mixture was then incubated at 37 °C for 36 h. The separation and purification of the released glycans from the were performed on tC18 RP cartridge, which was preconditioned with 2 mL of methanol and 2 mL of water. After loading the sample, the cartridge was washed twice with 200 µL of water. The flowthrough containing the released glycans, along with the column washes, was collected and dried under vacuum.

### Reduction of *N*-glycans and GSL glycans

The Glycans were reduced by adding 25 µL of 1 M NaBH_4_ (Sigma-Aldrich, Cat. no. 213462) in 100 mM potassium hydroxide (Sigma-Aldrich, Cat. no. 757551), through incubation at 50 ℃ for 3 h. Three microliters of glacial acetic acid (Merck, Cat. no. 5.33001.0050) was added to quench the reaction.

### Desalting of reduced glycans

Reduced *N*-, *O*-, and GSL glycans were desalted using cation exchange. The columns were self-packed with 25 µL of strong cation-exchange resin (50W-X8) slurry (BIO-RAD, cat. no. 1421451) in Methanol onto a ZipTip C_18_ resin tip. The columns were preconditioned sequentially with 3 × 50 µL of 1 M HCl (Sigma-Aldrich, cat. no. 320331), 3 × 50 µL of methanol, and 3 × 50 µL of ultrapure water. The samples were applied to the preconditioned columns, and the original sample tube was rinsed with 50 µL of ultrapure water, which was then added to the column. The glycans were subsequently eluted with 2 × 50 µL aliquots of ultrapure water. The eluted glycans were then dried and treated with 4 × 100 µL of methanol to remove the borate salt.

### PGC clean-up of *N*-, *O*- and *GSL* glycans

Carbon material was removed from the carbograph SPE cartridge (BGB Analytik AG, cat. no. SPU-5122423) and a slurry was prepared with a concentration of 50 mg/mL in methanol. About 25 µL of the slurry was loaded into a ZipTip with 0.6 µL C18 resin. The columns were pre-conditioned with 3 × 50 µL of 0.1% trifluoroacetic acid (Sigma-Aldrich, cat. no. 302031) and 3 × 50 µL of 90% acetonitrile (Sigma-Aldrich, cat. no. 1.00029), 0.1% trifluoroacetic acid. The glycan samples were loaded in 50 µL of 0.1% trifluoroacetic acid. Finally, the samples were eluted with 50 µL of 50% acetonitrile, 0.1% trifluoroacetic acid. The eluted glycans were then dried using a speed vac concentrator (Thermo Scientific; Savant SPD131DDA) and stored at −20 °C until further use.

### PGC-LC-ESI-MS/MS of released *N*- and *O*- and GSL glycans

The PGC-LC-ESI-MS/MS glycomics analysis was performed using an amaZon speed ion trap (IT-MS, Bruker, Bremen, Germany) linked to an Ultimate 3000 UHPLC system (Dionex/Thermo). Five microliters of glycan sample were injected by the autosampler and separated using an analytical PGC column (Hypercarb™ PGC Column, Thermo Fisher Scientific, 3 μm particle size, inner diameter 1 mm x 30 mm, maintained at 45 °C). Ten millimolar ammonium bicarbonate was used as buffer A and 70% acetonitrile in 10 mM ammonium bicarbonate was used as solvent B in the mobile phase. The flow rate was set at 15 μL/min for *N*-, *O*-, and GSL glycans. The gradient system used to separate the *N*-glycans was as follows: 1% B at 0 min; 14% B at 6 min; 25% B at 25 min; 70% B at 45 min; 98% B at 50 min; 1% B at 51 min; 1% B at 65 min. *O*- and GSL glycans were separated using the following gradient: 1% B at 0 min; 25% B at 25 min; 70% B at 45 min; 98% B at 47 min; 1% B at 51 min; 1% B at 65 min. Eluted glycans were then ionised using ESI in negative ion mode and detected with an ion-trap mass analyser. A *m/z* range of 460–1,800 was set for data-dependent precursor scanning. The collision induced dissociation (CID) fragmentation was used as the fragmentation method where the five most intense precursors of each MS scan were selected for tandem MS. Capillary voltage was set at 3.3 kV, with nitrogen drying gas flow rate 6 L/min at 8.70 psi nebulizer pressure. An ICC target of 70,000 and maximum accumulation time of 100 ms were used. Smart parameter setting (SPS) was set to 1200 *m/z*.

### Glycan structure determination and relative quantitation

The MS data was manually analysed using Compass Data analysis version 4.4 (Bruker Daltonics) and then quantitated with Skyline version 22.2^14^. Monosaccharide compositions were determined using the Glycomod tool available on the Expasy^15^ with a mass tolerance of 0.5 Da. The glycan structures were assigned manually from the tandem mass spectra and characterized with the GlycoWorkbench version 2.1^16^. The fragmentation spectra were also further matched with UniCarb KB to confirm all the structures^17^. Normalised relative intensities of individual glycan structures were calculated in Microsoft Excel and using GraphPad Prism, and to perform statistical analysis

### Statistical analysis

All the experiments were done in triplicates, and statistical analysis was performed in GraphPad Prism (version 9). *P* values were calculated by performing an unpaired two-tailed Welch’s *t*-test at 95% confidence level. Data visualization was performed in R Studio, with final figure preparation and refinement done in Adobe Illustrator.

### Glycan Representation

Glycan structures were visualized by cartoons generated using GlycoWorkbench Glycans were depicted according to the SNFG (Symbol Nomenclature for Glycans) notation: *N*-acetylglucosamine (*N*; blue square), fucose (F; red triangle), mannose (H; green circle), galactose (H; yellow circle), *N*-acetylneuraminic acid (S; purple diamond), and *N*-glycolylneuraminic acid (G; light blue diamond).

## Results

### HPIV-3 infection triggers significant changes in *N*-glycan expression in the host cell

We hypothesized that viral infection could alter glycan expression in host cells, which may influence processes such as progeny virion release, cell-cell and cell-matrix interactions, and overall cell signalling. To study the impact of infection on host-cell glycosylation, we analysed *N*-glycan expression of cell-extracted and secreted proteins from both mock-infected and HPIV-3 infected A549 cells, at 24 h and 48 h timepoints post-infection. In total, 70 *N*-glycan species (including structural and compositional isomers) were observed across infected and mock-infected cells, varying from 5 to 17 monosaccharides (Table S1). The glycan species were manually interpreted and validated based on the existence of specific fragment ions in their MS/MS spectra^18^. The identified *N*-glycans were grouped based on structural features (oligomannose, complex sialylated, complex neutral, hybrid, and paucimannose). Relative abundances of each glycan species from infected and mock-infected cells were statistically analysed (Table S2). The differential expression of *N*-glycans in infected and mock-infected cells is provided in Figure 1a-c.

**Figure 1:**
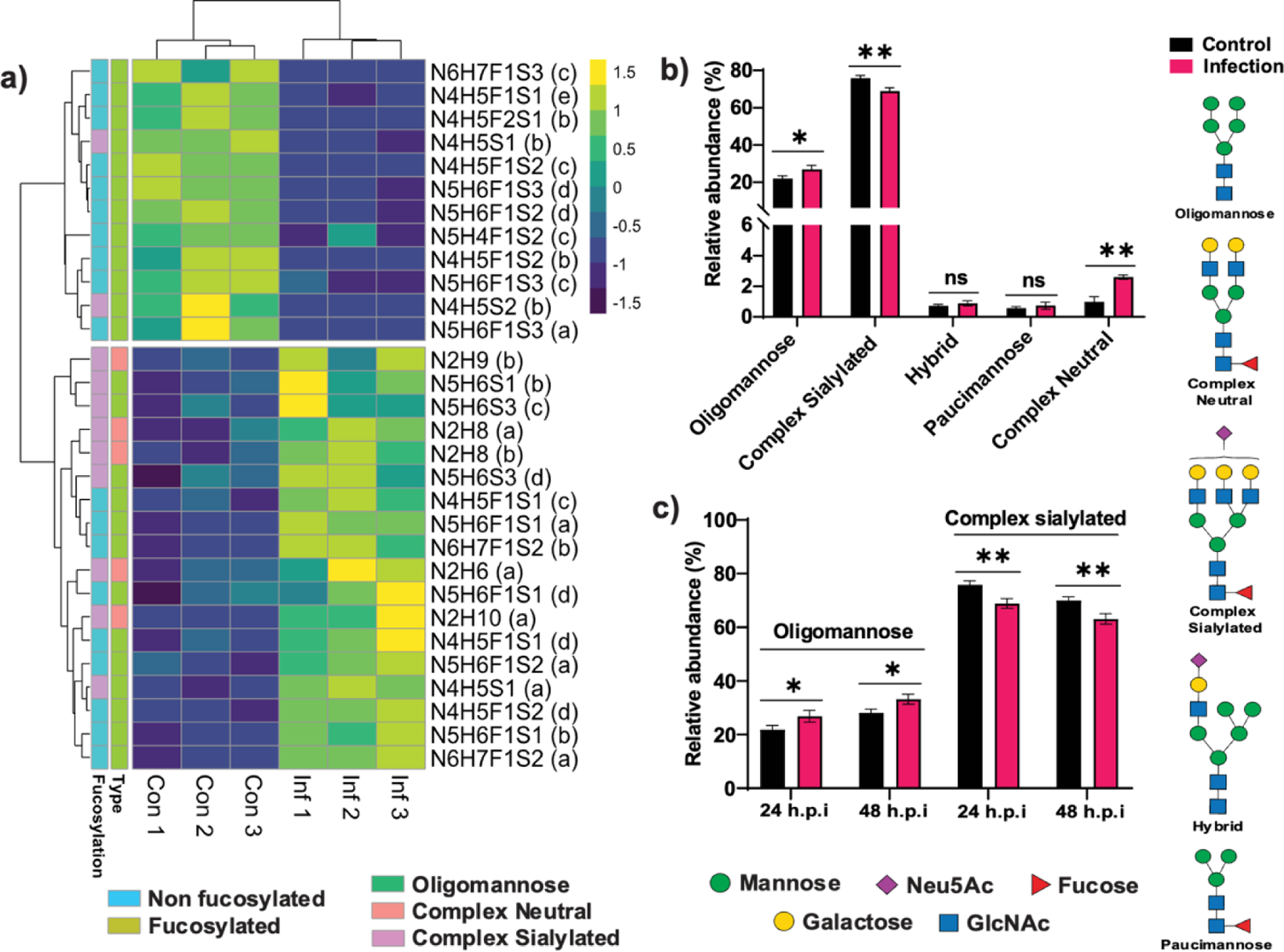
HPIV-3 infection modulates expression of *N*-glycans on cell-extracted proteins. a) Heatmap representing the relative abundance of significantly altered *N*-glycans before and 24 h post-infection. b) Differential expression of all major types of *N*-glycans on infected and mock-infected cells at 24 h post-infection. c) Altered expression of oligomannose-type and sialylated complex-type *N*-glycans at 24 h and 48 h timepoints post-infection. n=3, *P<0.05, **P<0.01, ns: not significant, unpaired Welch’s t-test. N: *N*-acetylhexosamine; H: Hexose; F: Fucose; S: *N*-acetylneuraminic Acid; h.p.i: hours post-infection.

In the context of total *N-*glycan composition, we observed that sialylated complex-type *N-*glycans and oligomannose-type glycans were the most abundant, comprising more than 90% of total *N*-glycans detected in both mock-infected and HPIV-3 infected A549 cells (Table S2). We observed a significantly higher expression of oligomannose-type *N*-glycans (from 21.87% ± 1.50% in mock-infected to 26.87% ± 2.18% in infected cells) at 24 h post-infection (Figure 1b). We also observed diverse sialylated structures ranging from mono-sialylated biantennary to tri-sialylated tri-antennary complex glycans (Table S1). HPIV-3 infection resulted in significant reduction of sialylated complex-type *N*-glycans in infected cells, decreasing from 75.85% ± 1.47% in mock-infected to 68.90% ± 1.81% in infected cells (Figure 1b). To identify if the altered expression of glycans in infected cells is stable, we analysed the glycan expression also at 48 h post-infection. Notably, similar changes in both oligomannose-type and sialylated complex-type *N*-glycans were also observed at 48 h post-infection. For example, an approximately 5% increase in oligomannose-type and a 7% decrease in sialylated complex-type *N*-glycans were detected in infected cells (Figure 1c). Moreover, we saw a subtle, yet statistically significant, increase (~1.5%) in the expression of complex neutral-type *N*-glycans upon 24 h and 48 h post-infection. Two other major glycan types, hybrid and paucimannose, remained unchanged during infection at 24 h and 48 h.

Cell secreted proteins play important roles in maintaining cell communication and signalling, and glycans that decorate these secreted proteins are key to these interactions^19^. Considering these important roles, we analysed the glycans released from secreted proteins of both infected and mock-infected cells and compared their relative abundances (Figure 2a).

**Figure 2:**
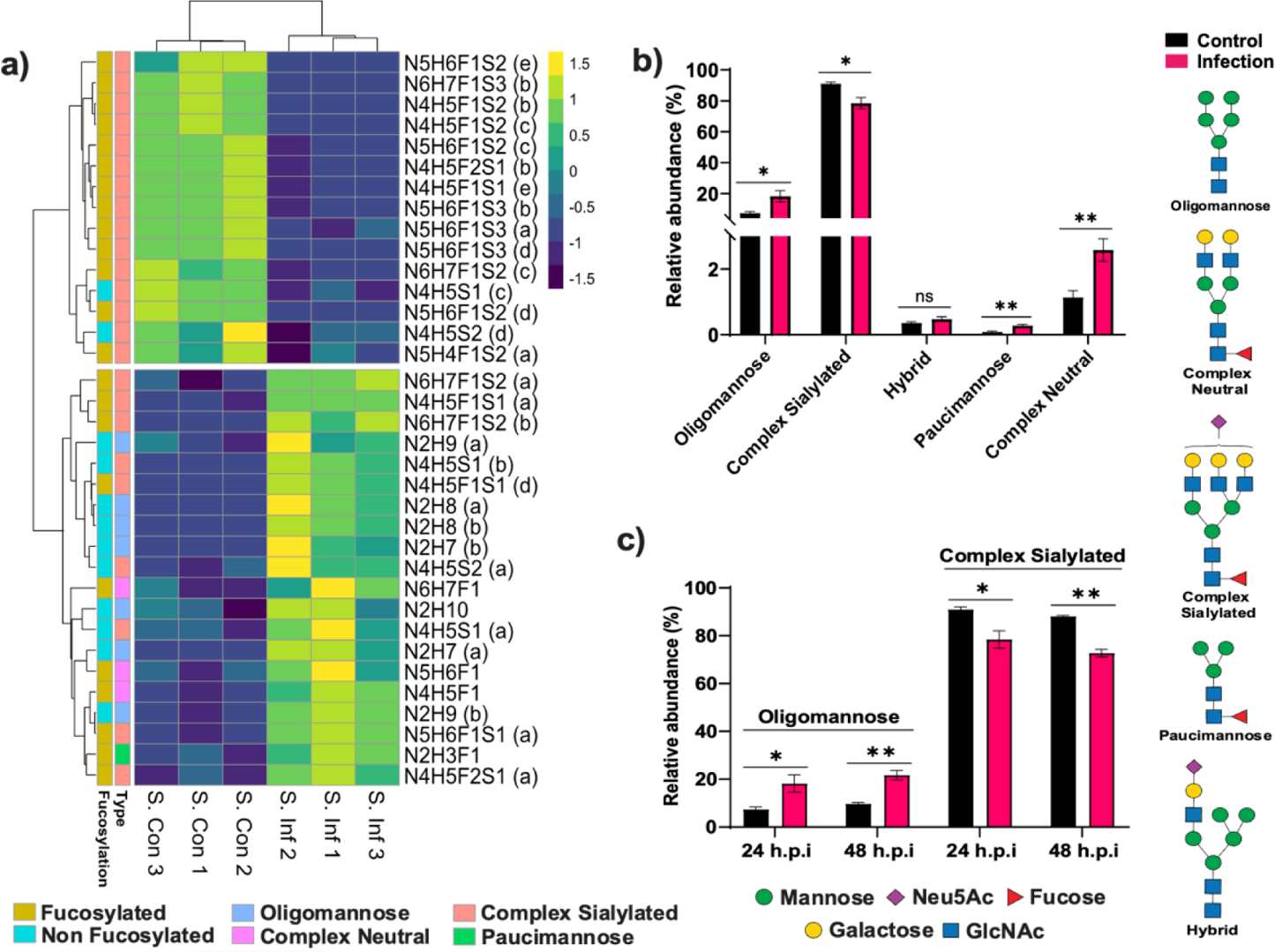
HPIV-3 infection modulates expression of *N*-glycans on secreted proteins. a) Heatmap represents the relative abundance of significantly altered *N*-glycans before and at 24 h post-infection. b) Differential expression of all major types of *N*-glycans in infected and mock-infected cells at 24 h post-infection. c) Altered expression of oligomannose-type and sialylated complex-type *N*-glycans at 24 h and 48 h post-infection. n=3, *P<0.05, **P<0.01, ns: not significant, unpaired Welch’s t-test. N: *N*-acetylhexosamine; H: Hexose; F: Fucose; S: *N*-acetylneuraminic Acid; h.p.i: hours post-infection.

The *N*-glycome composition of secreted proteins differed slightly from that of cell-extracted proteins. Approximately 90% of *N*-glycans on secreted proteins were processed from oligomannose to sialylated complex-type structures, whereas cell-extracted proteins contained about 70-74% sialylated complex-type *N*-glycans (Table S1). However, in agreement with *N*-glycans on cellular proteins, expression of sialylated complex-type glycans was decreased from 91% ± 1.08% in mock-infected control to 78.44% ± 3.66% in HPIV-3 infected cells (24 h post-infection). We also observed a significant increase of ~11% in oligomannose-type glycans in response to viral infection (24 h post-infection) (Figure 2b). Notably, these changes were nearly twice as pronounced as those observed in cell-extracted proteins, which showed a 7% decrease in sialylated complex-type and a 5% increase in oligomannose-type *N*-glycans. Likewise, HPIV-3 infection also modulated the *N*-glycome composition of secreted proteins at 48 h post-infection. The level of oligomannose-type glycans was augmented by more than 12%, whereas the expression of sialylated complex-type *N*-glycans was reduced by more than 15% on secreted proteins collected from HPIV-3 infected cells compared to mock-infected control (Figure 2c). The complex neutral type comprised of 1% to 2% of the total secreted protein *N*-glycome in mock-infected cells. However, the expression of this glycan type was observed to be doubled at both 24 h and 48 h post-infection (Figure 2b). There were no significant changes observed in relative intensities of hybrid and paucimannose-type *N*-glycans on secreted proteins after infection irrespective of the time points (Figure 2b).

### Differential expression of oligomannose and sialylated complex-type *N*-Glycans at the individual glycan level

Next, we examined whether changes in oligomannose and sialylated complex-type *N*-glycans followed a similar pattern at the level of individual glycan species. A detailed analysis enabled us to identify that, although overall abundance of oligomannose-type *N*-glycans increased from the mock-infected control (21.87% ± 1.50%) to infected cells (26.87% ± 2.18%), not all oligomannose species exhibited a similar trend. We observed that individual oligomannose species were either increased or remained constant post-infection (Figure S1, S2). Oligomannose glycans containing 5 to 9 mannose units were identified across infected and mock-infected cells. Of these, both isomers of N2H8, one isomer of N2H9, N2H6, N2H7 and N2H10 were increased at 24 h post-infection on cell-extracted proteins (Figure S1a). Likewise, each isomer of N2H7 to N2H9 was upregulated, whereas others remained constant after 48 h of infection when compared to the mock-infected control (Figure S2a). This trend was consistent for secreted proteins both at 24 h and 48 h post-infection, as at least one isomer from N2H6 to N2H9 was significantly elevated (Figure S1b & S2b). We also detected a range of sialylated structures, from mono-sialylated bi-antennary to tri-sialylated tri-antennary complex-type *N*-glycans, on both cell-extracted and secreted proteins. Using a PGC-LC-ESI-MS/MS workflow to enable the separation of isomers based on their corresponding retention time, we found that the majority of the sialylated structures displayed multiple isomeric forms (Table S1). Overall, there was an approximately 7% reduction in sialylated complex-type *N*-glycans in infected cells compared to their mock-infected counterparts on the cell-extracted proteins. However, this overall decrease did not apply uniformly to all sialylated complex-type *N*-glycan species, as some were downregulated while others were either upregulated or remained unchanged in infected cells (Figure S3 to S8). For example, singly sialylated bi-antennary structures with (N4H5F1S1) and without core fucose (N4H5S1), along with some doubly, triply sialylated bi-antennary (N4H5F1S2) and tri-antennary (N5H6S3) structures were found to be upregulated in infected cells (Figure S3). On the other hand, structural isomers of the singly sialylated bi-antennary structures with the composition of N4HS5F1S1, along with doubly and triply sialylated bi-antennary (N4H5F1S2) and tri-antennary (N5H6F1S3) *N*-glycans were downregulated upon HPIV-3 infection (Figure S4). These diverse changes in the expression of sialylated complex-type *N*-glycans were detected across cell-extracted and secreted proteins, regardless of post-infection time points (Figure S5-S10). Together, these findings highlight that HPIV-3 infection results in selective remodelling of both oligomannose and sialylated complex-type *N*-glycans, with distinct changes observed at the level of individual glycan species.

### HPIV-3 infection modulates expression of sialic acid isomers on *N*-glycans

HPIV-3 binds to sialylated glyco-epitopes on host cells via its HN protein to initiate infection and later uses the same protein’s neuraminidase activity to cleave sialic acids (Neu5Ac), enabling the release of newly formed virions^20^. Considering this, we hypothesize that HPIV-3 infection could directly affect the expressions of glycans terminating in Neu5Ac. Structural isomers of sialylated complex-type *N*-glycan species arise from the linkage of Neu5Ac to the terminal galactose via either an α2,3 or α2,6 glycosidic bonds on each antenna of the glycan. Previous studies suggest that α2,3- and α2,6-linked Neu5Ac residues on galactose serve as receptors for HPIV-3, with the viral HN neuraminidase showing a preference for α2,3-linked Neu5Ac-containing glycans over their α2,6-linked counterparts^20^. Such differential recognition and cleavage specificity suggest that HPIV-3 may selectively modulate sialylation patterns during infection. Although sialic acid linkage isomers can be definitively characterized using linkage-specific neuraminidases, the linkage of structurally simple glycans bearing a single sialic acid residue can also be inferred based on their retention behaviour on PGC column, as demonstrated in previously published studies^21,22^. We observed three potential isomers of bi-antennary complex *N*-glycans with *m/z* 965.8^2-^ (N4H5S1). In agreement with previous studies, the first two structures eluting closely on the PGC column are arm isomers of an α2,6-linked mono-sialylated complex-type *N*-glycan, whereas the third structure, eluting 6 minutes later than the first two, represents an α2,3-linked isomer of the same composition (N4H5S1)^22^. Comparing the relative abundance of these species, we found that HPIV-3 infection results in a significantly lower expression of the α2,3-linked isomer in both cell-extracted and secreted proteins, irrespective of post-infection time points (Figure 3a & 3b). On the other hand, the α2,6-linked isomer of the same *N*-glycan composition was increased in infected cells when compared to the mock-infected control (Figure 3c & 3d).

**Figure 3:**
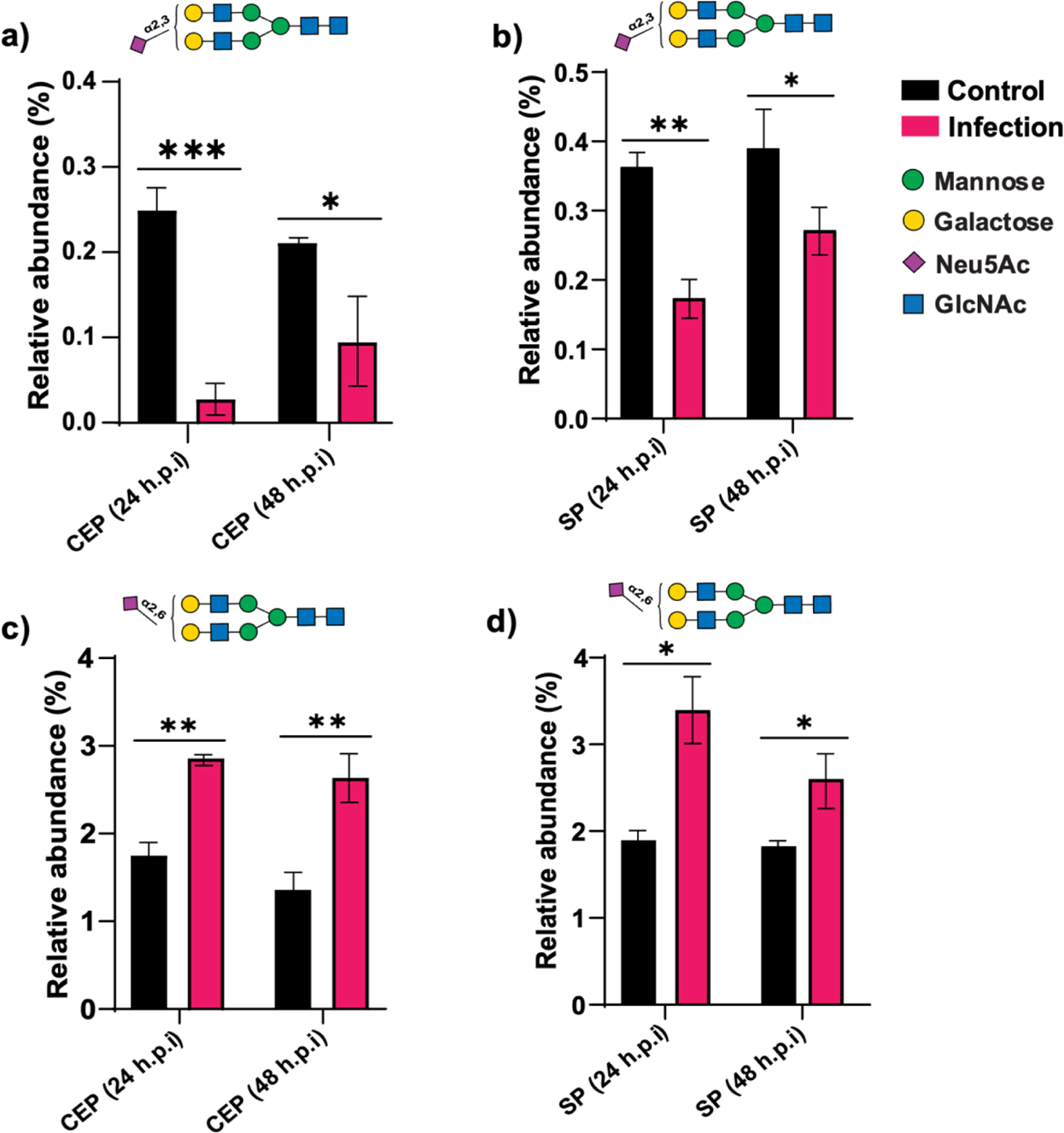
Differential expression of α2,3- and α2,6-linked isomers of mono-sialylated complex-type *N*-glycan (N4H2S1) in infected vs. mock-infected cells: (a) α2,3 on cell-extracted proteins, (b) α2,3 on secreted proteins, (c) α2,6 on cell-extracted proteins, (d) α2,6 on secreted proteins at 24 h and 48 h post-infection. n=3, *P<0.05, **P<0.01, ***P<0.001, unpaired Welch’s t-test. CEP: Cell extracted proteins; SP: Secreted proteins. h.p.i: hours post-infection.

### Alterations in *O*-glycan profiles induced by HPIV-3 infection

Since we identified significant modulation of the host-cell *N*-glycome upon HPIV-3 infection, we pondered if the *O*-glycome of host cell could also be altered. Like *N*-glycans, *O*-glycans also have a wide range of roles in the viral replication cycle, such as facilitating viral entry by acting as co-receptors and regulating viral navigation at the cell surface to assist virion progeny egress^23^. Changes in the *O*-glycan profile because of HPIV-3 infection could significantly affect the diverse and multifaceted functions mediated by *O*-glycans. To address this open question, we investigated *O*-glycan expression at 24 h and 48 h timepoints post-infection in both cell-extracted and secreted proteins. The profiling of *O*-glycans was carried out essentially as described for the *N*-glycans. In total, 9 *O*-glycans structures including structural isomers were detected in cell-extracted proteins from infected and mock-infected A549 cells (Table S1). The *O*-glycome of A549 cell is comprised of core 1 and core 2 type glycans with core 2 type the most abundant (from 67% to 93% across mock-infected and infected cells) (Table S2). A distinctive *O*-glycan profile was found on infected cell-extracted proteins when compared to their mock-infected counterpart (Figure 4).

**Figure 4:**
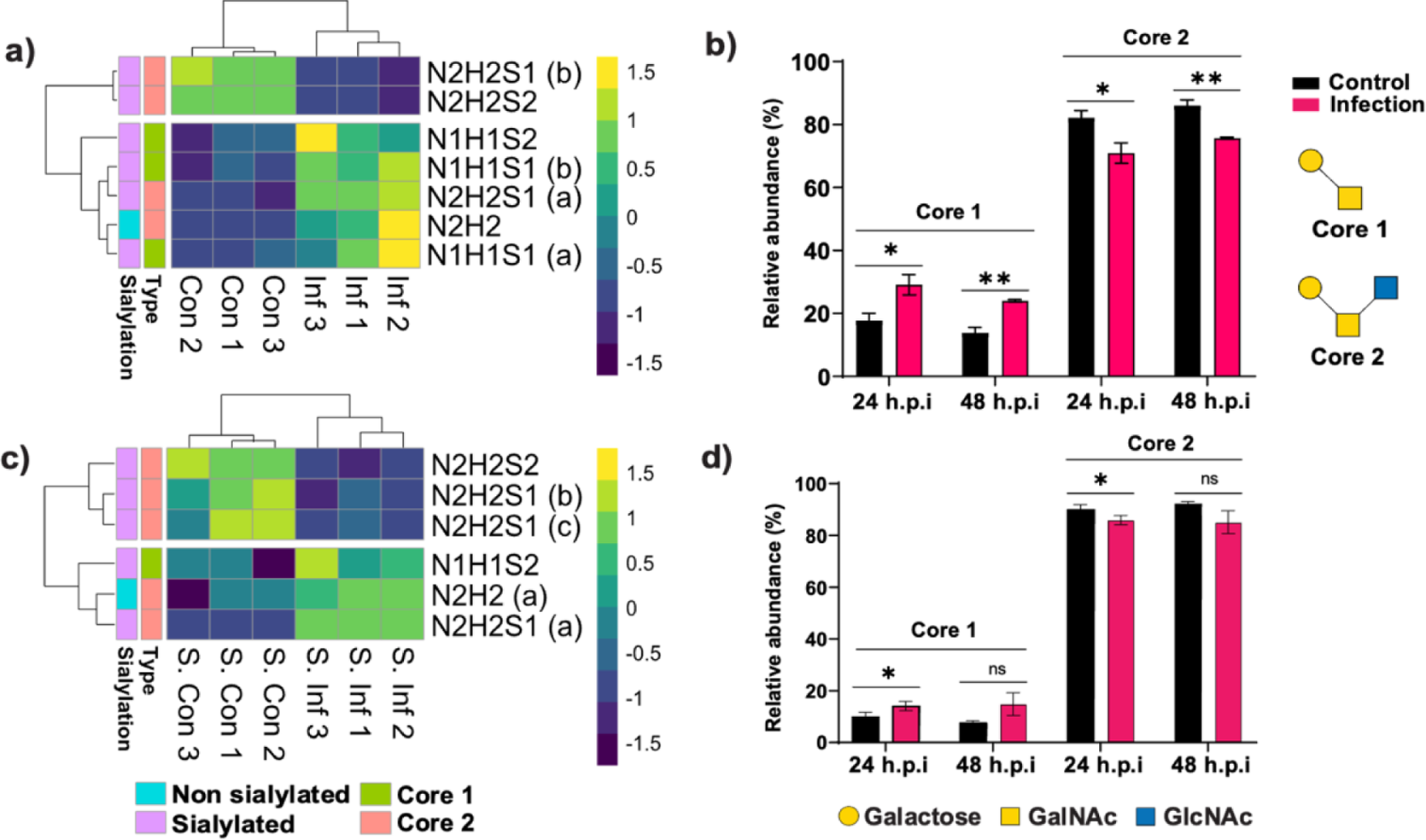
HPIV-3 infection modulates expression of *O*-glycans on cell-extracted and secreted proteins. a) Heatmap represents the relative abundance of significantly altered *O*-glycan species on cell-extracted proteins before and 24 h after infection. b) Differential expression of two major core types on cell-extracted proteins at 24 h and 48 h post-infection. c) Heatmap represents the significantly altered *O*-glycan species on secreted proteins before and 24 h post-infection. d) Differential expression of two major core types on secreted proteins at 24 h and 48 h post-infection. n=3, *P<0.05, **P<0.01, ***P<0.001, ns: not significant, unpaired Welch’s t-test. N: *N*-acetylhexosamine; H: Hexose; F: Fucose; S: *N*-acetylneuraminic Acid; h.p.i: hours post-infection.

We observed that core 2 type *O*-glycans from host cell-extracted proteins were downregulated upon HPIV-3 infection at 24 h post-infection timepoint, reducing from 82.17% ± 2.20% in mock-infected cells to 70.90% ± 3.23% in infected cells (Figure 4b). Moreover, the downregulation of the core 2 type was compensated by an approximately 12% increase of core 1 type in infected cells. A similar response was observed at 48 h post-infection. Next, we sought to determine if the expression of *O-*glycans on secreted protein are also altered upon HPIV-3 infection. A moderate decrease, from 90.10% ± 1.78% in mock-infected to 85.91% ± 1.78% in infected cells, of secreted protein core 2 type glycans and a corresponding increase, from 9.93% ± 1.78% in mock-infected to 14.10% ± 1.77% in infected cells, of secreted protein core 1 type glycans was found at 24 h timepoint post-infection (Figure 4d). The same trend was found at 48 h timepoint post-infection (Figure 4d).

Interestingly, upon investigating individual *O*-glycan species, we observed a similar pattern of infection-induced alterations across cell-extracted and secreted proteins irrespective of post-infection time points. For example, the expression of the di-sialylated core 2 species on cell-extracted proteins, characterized by an *m/z* of 665.2^2-^ (composed of N2H2S2), was reduced by approximately three-fold, with a ~27% and 47% decrease at the 24 h and 48 h timepoints post-infection, respectively in infected cells (Figure 5a and 5b). In contrast, the mono-sialylated core 2 glycan [N2H2S1(a)] showed a remarkable increase in expression of ~26% and 37% at 24 h and 48 h timepoints post-infection, respectively upon infection (Figure 5a and 5b).

**Figure 5:**
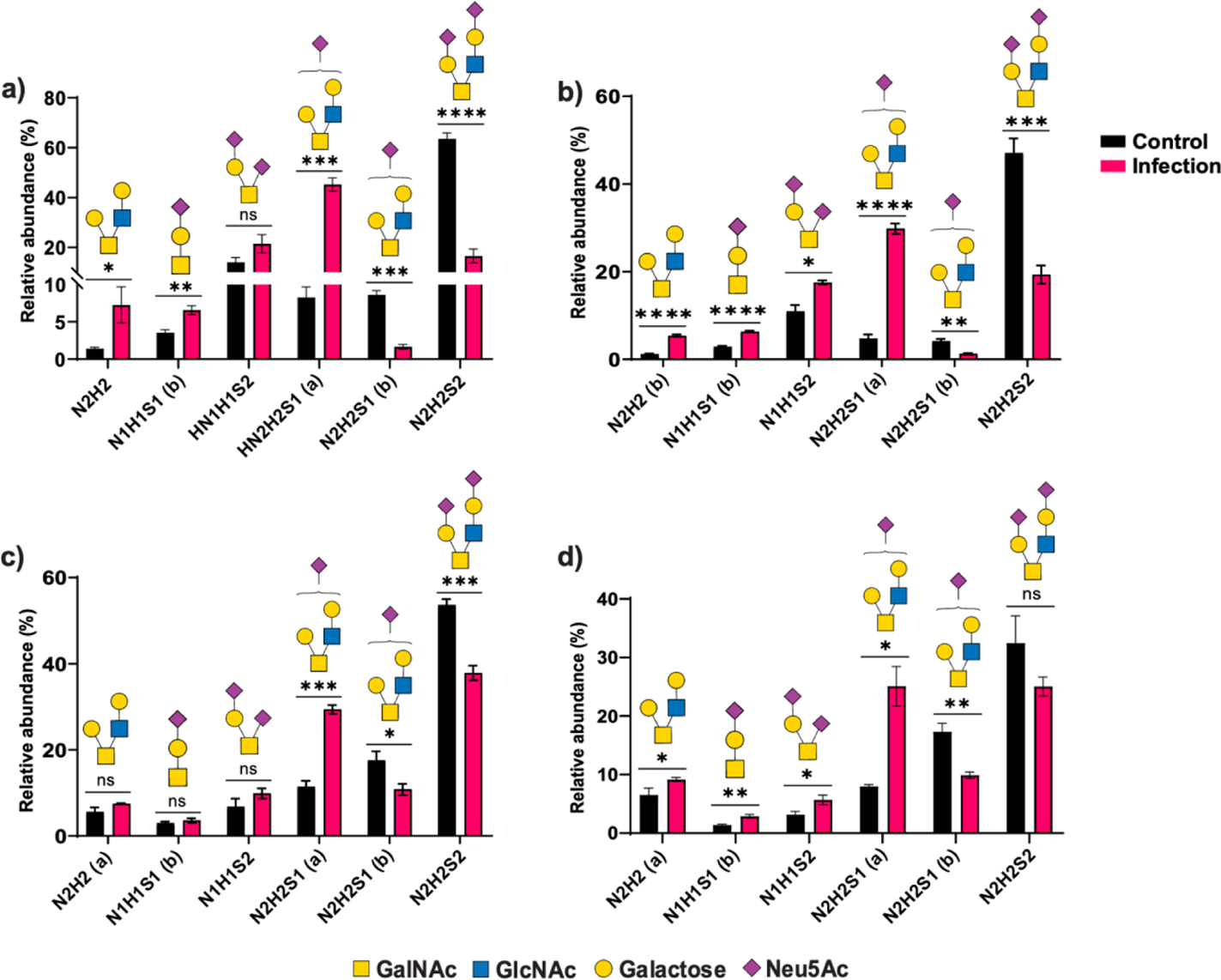
HPIV-3 triggers modulated *O*-glycan expression in infected and mock-infected cells. Relative abundance of all the significantly altered *O*-glycan species on cell-extracted proteins at a) 24 h, and b) 48 h post-infection. Relative abundance of all the significantly altered *O*-glycan species on secreted protein at c) 24 h, and d) 48 h post-infection. n=3, *P<0.05, **P<0.01, ***P<0.001, ns: not significant, unpaired Welch’s t-test. N: *N*-acetylhexosamine; H: Hexose; S: *N*-acetylneuraminic Acid.

This trend was similarly observed for secreted proteins, independent of the post-infection timepoint (Figure 5c and 5d). Moreover, a structural isomer of the same composition [N2H2S1(b)] was reduced upon infection across cell-extracted and secreted proteins (Figure 5a-d). Interestingly, a core 1 type doubly-sialylated *O*-glycan, with a composition of N1H1S2, was found to be significantly elevated in infected cells for both cell-extracted and secreted proteins at each post-infection timepoint (Figure 5a-d). We observed a slightly favourable expression of core 1 type *O*-glycans compared to core 2 type, along with altered expression of glycans carrying specific sialylated glyco-motifs in response to infection. These changes were consistent for glycans released from both cell-extracted and secreted proteins, regardless of post-infection time points.

### HPIV-3 infection induces remodelling of glycosphingolipid (GSL)-associated glycans

GSL glycans represent a significant portion of the host glycome and are primarily located on the cellular plasma membrane, where their sugar moieties are exposed to the external environment^14^. These glycans are involved in various biological processes, including infections^24^. Some viruses (e.g HPIV-3^20^, HIV^25^, Polyomavirus^26^) are historically known to utilize GSL glycans in their life cycle. However, no studies have reported host cell GSL glycan dynamics in response to parainfluenza viral infection. To characterize GSL glycans, GSLs were extracted from both infected and mock-infected cells before the enzymatic release of the glycans from the head groups, which were then analysed by PGC-LC-ESI-MS/MS. In total, 12 GSL glycan species were detected, and the relative intensities of each glycan species were compared between infected and mock-infected control (Table S1). The structure of some of the GSL glycans could only be partially resolved in this study due to the absence of informative fragment ions in tandem MS, and the lack of *exo*-glycosidase digestion in our study. Therefore, there were no GSL glycans class wide comparison made between infected and mock-infected controls. However, major alterations have been detected at the individual glycan level in infected cells (Figure 6).

**Figure 6:**
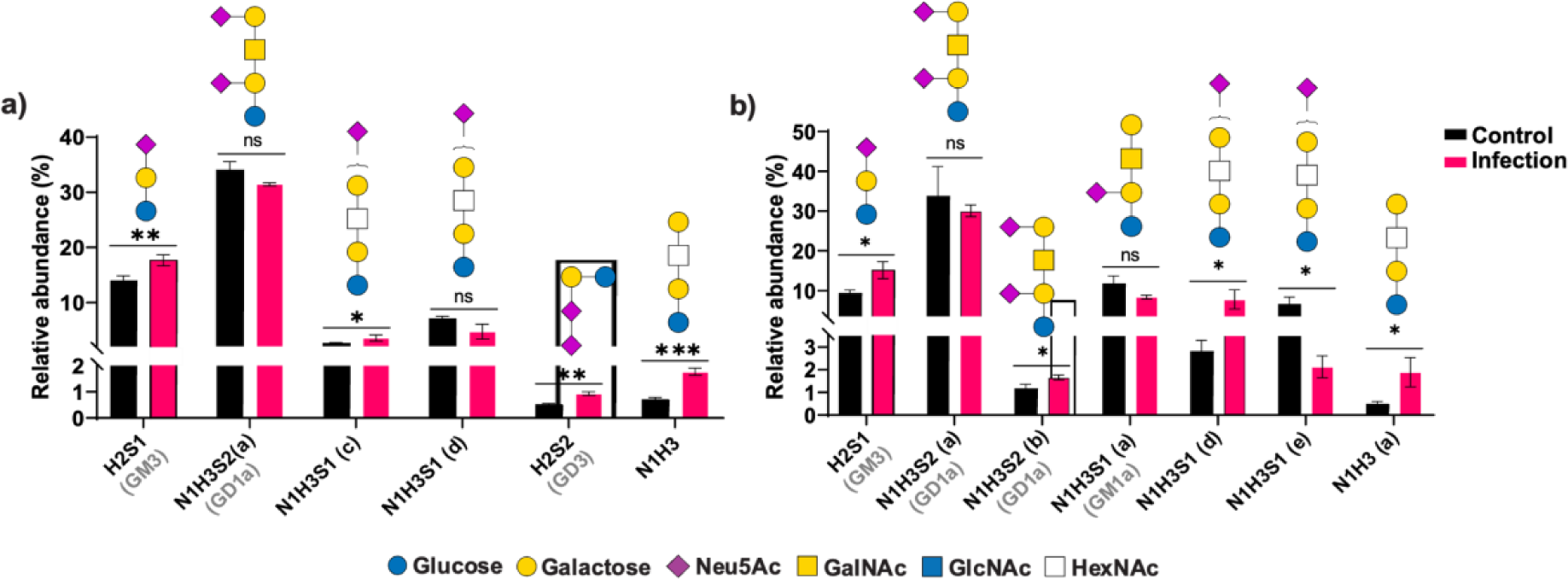
HPIV-3 infection triggers altered expression of GSL glycans. Relative expression of all the significantly altered GSL glycan species at a) 24 h, and b) 48 h post-infection. n=3, *P<0.05, **P<0.01, ***P<0.001, ns: not significant, unpaired Welch’s t-test. N: *N*-acetylhexosamine; H: Hexose; S: *N*-acetylneuraminic Acid.

Gangliosides containing one and two Neu5Ac residues were the most abundant GSL glycan types in A549 cells followed by the neo(lacto)-series. The GSL glycan, comprised of H2S1 (GM3), appeared to be increased in expression from 14.0% ± 0.84% in mock-infected cells to 17.66% ± 1.01% in infected cells, 24 h post-infection (Figure 6a). The increased expression of GM3 remained stable even at 48 h timepoint post-infection (Figure 6b). Another ganglioside comprised of N1H3S2 (GD1a), the most abundant GSL glycan in A549 cells, was reduced in expression from 34.02% ± 1.57% in mock-infected cells to 31.44% ± 0.27% in infected cells, at 24 h timepoint post-infection. A similar trend was observed at 48 h timepoint post-infection despite the change not being statistically significant. Additionally, several other GSL glycan species were also observed to be altered in response to HPIV-3 infection. Two partially resolved isomers with the composition of H3N1S1 (presumably sialylated neo(lacto)-series or GM1 gangliosides) showed an opposite trend to each other upon infection. The changes were more prominent at 48 h timepoint post-infection when compared to the mock-infected control. Interestingly, a previous study reported that the sialylated neo(lacto)-series, with the same composition, is recognised by HPIV-3 using a solid-phase binding assay^20^. Higher expression of one of the H3N1S1 isomers in infected cells could be attributed to the reduced expression of GD1 (H3N1S2), as HPIV-3 HN mediated sialic acid cleavage from GD1 glycan would potentially generate a glycan with the composition of H3N1S1, thereby contributing to overall increase of that species upon infection. Furthermore, the expression of GD3 (H2S2) was found to be elevated only at 24 h timepoint post-infection and no significant change was observed at 48 h timepoint post-infection. A precursor glycan, belonging to either gangliosides or neo(lacto)-series, with composition of H3N1 was found to be elevated at both 24 h and 48 h timepoints post-infection.

## Discussion

Every cell in nature is decorated with a dense layer of carbohydrates (glycans) covalently linked to proteins or lipids, forming the glycocalyx. These glycans form the foundation of a universal biological language known as the glycome, playing active roles in numerous cellular processes^27^. However, microorganisms such as viruses exploit these cell surface resident glycans by using them as receptors or co-receptors^28^. Additionally, viruses take advantage of the host-cell glyco-machinery to decorate their own surface proteins with glycans, acting as protective shields to evade the host immune response^29,30^. During human evolution, viruses along with other pathogens such as bacteria and parasites have co-evolved with their human host and consequently have actively participated in the remodelling of the human cell glycome^31^. For instance, it is widely accepted that evolutionary pressure from pathogens played a significant role in the loss of certain carbohydrates, such as *N*-glycolylneuraminic acid (Neu5Gc) and α-Gal, in human^32,33^.

To date, only a few studies have described the changes in host-cell glycosylation upon viral infection and the role of an altered glycome expression in triggering innate immune response^6,34,35^. For example, a lectin microarray based study has suggested a significant increase of oligomannose-type glycans in ferret lung upon influenza virus infection, which was further found to be associated with disease severity^6^. More recently, several studies on SARS-CoV-2 infection observed an altered glycosylation profile on antibodies against the viral spike protein with lower fucosylation and sialylation, that ultimately affects the effector functions of antibodies^34,35^. However, these studies either did not employ more sensitive techniques like mass spectrometry or focused solely on the glycosylation of antibody (IgG) against viral protein, rather than examining the glycosylation of infected host-cells. To the best of our knowledge, there is no reported study on the host-cell glycome response to HPIV-3 infection. Thus, we employed a highly sensitive mass spectrometry technique coupled with porous graphitized carbon liquid chromatography to reveal the global glycosylation (*N-*, *O*- and GSL glycans) changes of HPIV-3-infected cells. Our analysis revealed an approximately 5% to 12% increase of oligomannose-type *N*-glycans on cell-extracted and secreted proteins, both at 24 h and 48 h post-infection timepoints. This increase may reflect a host cellular response to infection-induced activation of the unfolded protein response (UPR), as has been proposed in other viral systems, such as influenza infection in the ferret lung^6^. The UPR is a well-characterized cellular stress mechanism that is activated when unfolded or misfolded proteins accumulate in the endoplasmic reticulum (ER), disrupting protein homeostasis^36^. In response, the cell initiates UPR signalling to restore ER function by enhancing the folding capacity of the ER, reducing protein translation, and promoting the degradation of misfolded proteins^36^. One important hallmark of UPR activation is the increase in oligomannose-type *N*-glycans, which are essential intermediates in the early stages of protein glycosylation and folding^6,37^. These glycans are particularly important for the quality control of nascent glycoproteins, as they interact with ER-resident chaperones such as calnexin and calreticulin^37^. This interaction ensures that only properly folded proteins proceed along the secretory pathway, while misfolded proteins are retained and targeted for ER-associated degradation (ERAD)^32^. In the context of HPIV-3 infection, we observe an increase in oligomannose-type glycans which may reflect a host-cellular response to the stress placed on the ER to fold both viral and cellular proteins. Viral infection frequently disrupts ER homeostasis, not only due to synthesis of viral proteins but also through inflammation-induced oxidative stress and altered calcium signalling^38,39^. As a result, the host cell may ramp up UPR signalling to maintain protein homeostasis, not only for its own proteins such as membrane receptors, transporters, or immune-modulatory proteins, which are critical for cellular survival and antiviral responses, but also for the proper folding and processing of viral proteins^40,41^. Together, our findings support the hypothesis that the increased abundance of oligomannose-type *N*-glycans is driven by UPR-mediated adaptation aimed at preserving ER function and protein quality control during HPIV-3 infection. However, direct evidence for UPR activation in our system is still lacking. Future studies employing transcriptomics or proteomics analysis, particularly monitoring UPR markers such as BiP/GRP78, CHOP, and XBP1 splicing, will be essential to confirm whether the UPR pathway is indeed engaged in HPIV-3-infected cells and driving the observed glycomic changes^6^. Furthermore, the increased expression of oligomannose-type *N*-glycan was accompanied by overall reduction of sialylated complex-type glycans on cell-extracted and secreted proteins, irrespective of post-infection time points. We hypothesize that the elevated expression of oligomannose-type *N*-glycans contributes to the downregulation of sialylated complex-type *N*-glycans through two potential mechanisms. First, the accumulation of oligomannose structures may saturate the glycosylation machinery within the Golgi apparatus, thereby limiting the processing capacity for downstream glycan maturation. Second, as the biosynthesis of complex-type *N*-glycans requires the prior trimming of oligomannose precursors, excessive levels of these unprocessed glycans may reduce the availability of appropriately processed substrates necessary for complex glycan formation^5^. Future studies incorporating transcriptomic and proteomic analysis will benefit from evaluating the expression levels of Golgi-resident mannosidases and glycosyltransferases to confirm whether elevated oligomannose levels contribute to the reduction in complex-type *N*-glycan expression^6,42^. These studies, together with the existing glycomic data, may determine whether the reduction in complex-type *N-*glycan expression upon HPIV-3 infection is caused by limited substrate availability, as complex glycan synthesis requires the prior trimming of oligomannose, or by saturation of the glycosylation machinery due to an overload of oligomannose substrates.

Moreover, we were able to identify α2,3- and α2,6-linked isomers with the composition of N4H5S1, based on chromatographic separation, together with MS fragmentation^43^. Interestingly, HPIV-3 infection leads to a lower expression of *N-*glycans containing α2,3-linked sialic acid and an increase of its α2,6-linked counterpart, with the same composition (N4H5S1). A previous study found that HPIV-3 utilizes α2,3- and α2,6-linked *N*-acetylneuraminic acid (Neu5Ac) to galactose as receptors, with a higher HPIV-3 HN neuraminidase activity observed for α2,3-linked Neu5Ac^20^. Thus, the downregulation of α2,3-linked sialylated isomeric glycans, in this instance, could be attributed to the HPIV-3 HN neuraminidase activity, mediating the cleavage of receptor glycans at the end of each viral replication cycle^44^. Furthermore, we hypothesize that higher expression of α2,6-linked isomer of N4H5S1 in infected cells could be a consequence of the α2,3-linked sialic acid trimming of N4H5S2 (Figure S4, S6, S8, S10), as the HPIV-3 HN-mediated cleavage of N4H5S2 glycan would potentially generate glycan with the composition of HN4H5N1. To confirm the intrinsic role of HN protein on the reduction of sialylated *N*-glycans it would be interesting to treat infected cells with a specific inhibitor to HN protein^45^. Moreover, as glycan biosynthesis is a sequential enzymatic process, the modulation of sialylated complex-type *N*-glycans expression could also be the result of virus infection-mediated changes in enzyme expression related to glycan biosynthesis^27^. A bulk RNA-seq analysis of infected and mock-infected cells might enable us to identify the expression profiles of enzymes involved in glycan biosynthesis, thereby uncovering potential mechanisms underlying the observed changes in *N*-glycans upon infection.

The impact of HPIV-3 infection in the host-cell *O*-glycome is significant and we have observed that there is a significant reduction of core 2 type as well as a higher expression of core 1 type *O*-glycans across host cell-extracted and secreted proteins. A recent study also described an approximately 2.5-fold downregulation of core 2 *N*-acetylglucosamine transferase (GlcNAcT) enzyme upon HPIV-3 infection on A549 cells^42^. We propose that the reduction of core 2 type *O*-glycans could be the result of downregulation of the core 2 β1,6-*N*-acetylglucosaminyltransferase (Core 2 GlcNAc-T) enzyme that acts on core 1 type to make core 2 type *O*-glycans. Furthermore, concomitant with the observed down regulation of core 2 type *O*-glycans upon HPIV-3 infection, we also detected a down-regulation of doubly sialylated core 2 type glycans with a corresponding higher expression of singly sialylated core 2 type structures in infected cells. Similar for the downregulation of α2,3-linked sialylated isomeric *N*-glycans, we infer that the reduction of doubly sialylated core 2 type structures is caused by the neuraminidase activity of the HPIV-3 HN protein which subsequently results in the upregulation of similar glycans with one less Neu5Ac^46^. Again, to verify the specific role of the HPIV-3 HN protein in the reduction of doubly-sialylated *O*-glycans, it would be valuable to treat infected cells with a specific inhibitor of the HN protein^45^ and determine the expression of doubly-sialylated *O*-glycans. Nonetheless, these changes may have profound implications for host-cell glycosylation profiles, influencing immune responses and potentially contributing to the pathogenesis of the virus^6,42^.

GSL glycans significantly contribute to the formation of cellular glycome and some viruses including HPIV-3 are known to exploit GSL glycans in their life cycle^20^. In our analysis, gangliosides account for approximately 60% of the total GSL glycan population in A549 cells, underscoring their dominance in the glycan landscape of the respiratory epithelium. HPIV-3 triggers altered expression of gangliosides glycans including GM1, GM3, and GD3, which are often implicated in viral replication cycle^24^. These gangliosides have been previously implicated in modulating viral replication, either by serving as entry cofactors or by influencing intracellular signalling pathways essential for viral propagation^47,48^. For example, GM1 serves as entry receptors for simian virus 40 (SV40)^49^, while GM3 plays a dual role in cell physiology^50^ and viral pathogenesis, including facilitating influenza virus infection by modulating membrane dynamics and virus-host interactions^51^. Furthermore, GM3 has been shown to regulate cell proliferation and differentiation by modulating receptor tyrosine kinases such as epidermal growth factor receptor (EGFR)^52^. During viral infection, these signalling pathways are often hijacked to support viral replication and suppress host antiviral responses^53,54^. GD3 ganglioside is also implicated in viral infection, particularly those affecting nervous system^55^. Altogether, changes in ganglioside expression during HPIV-3 infection might represent either a viral adaptation to enhance conditions for its replication, or a host-cell mechanism to manage the infection.

The second most common type of GSL glycans detected in A549 cells was the neo(lacto)-series. A previous study reported a sialylated neo(lacto)-series glycan as a potential receptor for HPIV-3^20,46^. In our study we found a potential sialylated neo(lacto)-series glycan was downregulated upon HPIV-3 infection. We hypothesize that the cleavage of sialic acid, mediated by HPIV-3 HN, may contribute to the reduction of sialylated neo(lacto)-series glycans on HPIV-3 infected cells. Furthermore, few other glycans, which were not recognised as binding factors, have also been found to be altered after HPIV-3 infection in our study. We hypothesize a similar receptor specificity analysis using altered GSL glycans (GM1, GM3, GD3) could potentially identify few other GSL glycans as binding factors.

## Conclusion

Using advanced mass spectrometry-based glycomics, we investigated HPIV-3–induced alterations in the glycome of A549 human lung epithelial cells. Our analysis revealed approximately 116 distinct glycan species in both HPIV-3–infected and mock-infected cells, highlighting the complexity and diversity of the host-cell glycome. Importantly, this study provides the first experimental evidence that HPIV-3 infection induces marked alterations in the host cell glycome, spanning all three major glycan types (*N*-, *O*-, and glycosphingolipid glycans). These alterations suggest that HPIV-3 infection significantly remodels the host cell surface and secretory glycan landscape, potentially influencing key biological processes such as virus-host interactions, immune recognition, and intracellular signalling. Furthermore, our analytical approach provides a valuable platform for characterizing virus-induced glycomic changes and can be readily adapted to study other respiratory viruses, e.g., influenza virus, thereby paving the way for a better understanding of the roles of glycans in viral pathogenesis.

## Acknowledgement

This research was partially funded by Griffith University International Postgraduate Research Scholarship (GUIPRS) and Griffith University Postgraduate Research Scholarship (GUPRS) (P.K.D). We gratefully acknowledge funding from the National Health and Medical Research Council (NHMRC) (MvI., APP1047824 & GNT1196520) and a Peter Doherty Biomedical Early Career Fellowship (L.D., GNT1157150). We thank Dr. Thomas Litfin and Anuk Indraratna for assisting in data visualization.

## Author contributions

Conceptualisation: P.K.D., L.D., B.B., P.G., A.E.D, M.v.I.; methodology: P.K.D., P.G., A.E.D.; investigation: P.K.D.; analysis: P.K.D.; writing—original draft: P.K.D.; writing— review and editing: L.D., P.G., B.B., A.E.D., M.v.I.; funding acquisition: L.D., M.v.I.

## Abbreviations

HPIV: Human parainfluenza virus
HN: haemagglutinin-neuraminidase
H.p.i: Hour’s post-infection
CEP: Cell extracted protein
SP: Secreted protein
HN: *N*-acetylglucosamine
H: Hexose
dH: Deoxyhexose
Man: Mannose
N: *N*-acetylneuraminic Acid
G: *N*-glycolylneuraminic Acid
GSL: Glycosphingolipids
PGC: Porous graphitized carbon
ESI: Electrospray ionization
CID: Collision induced dissociation

## Supplementary Figures

**Fig S1.**
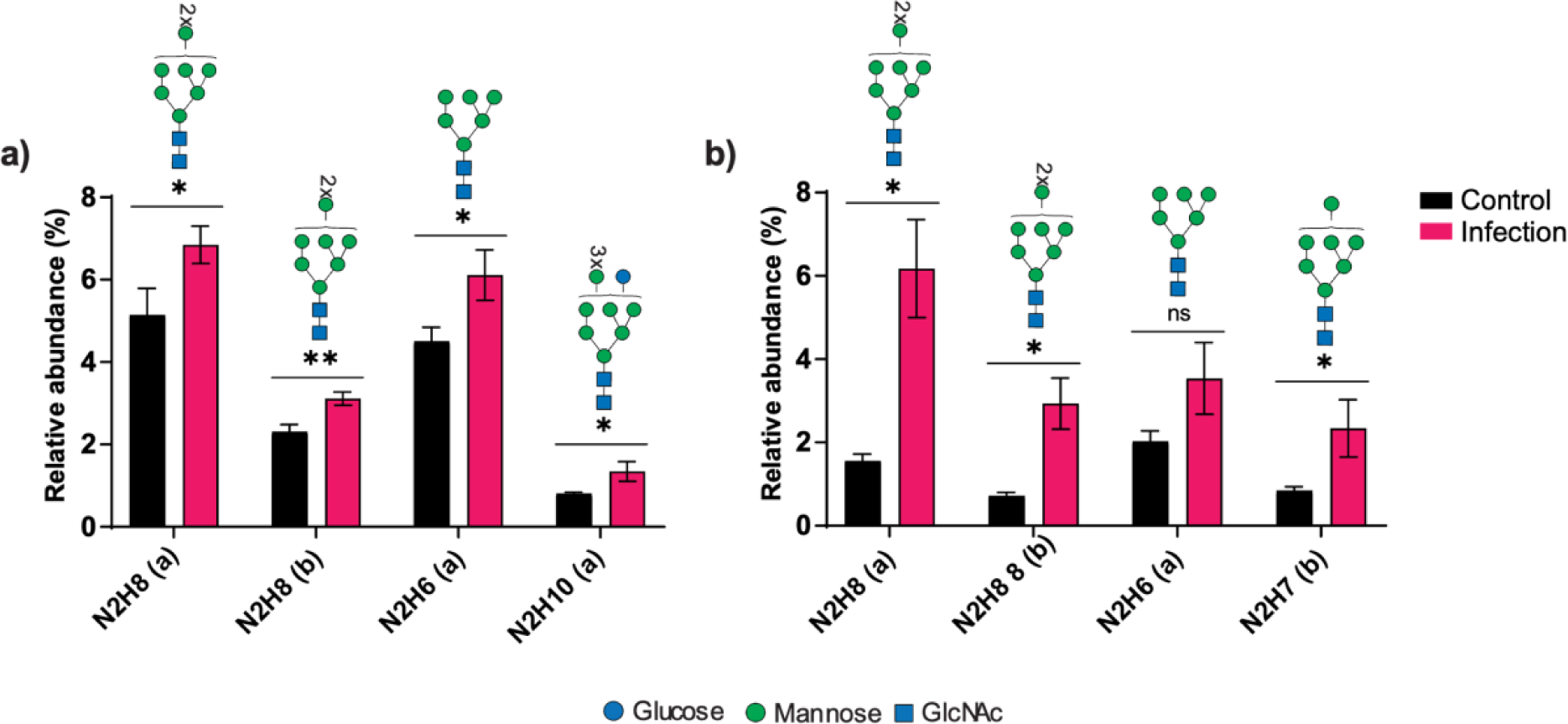
Alterations of oligomannose-type *N*-glycan on mock-infected and infected cells at 24 h post-infection. Relative abundance of significantly altered oligomannose type glycans on a) cell extracted proteins, b) secreted proteins. Man: Mannose; Glc: Glucose. n=3, *P<0.05, **P<0.01, ns: not significant, unpaired Welch’s t-test.

**Fig S2:**
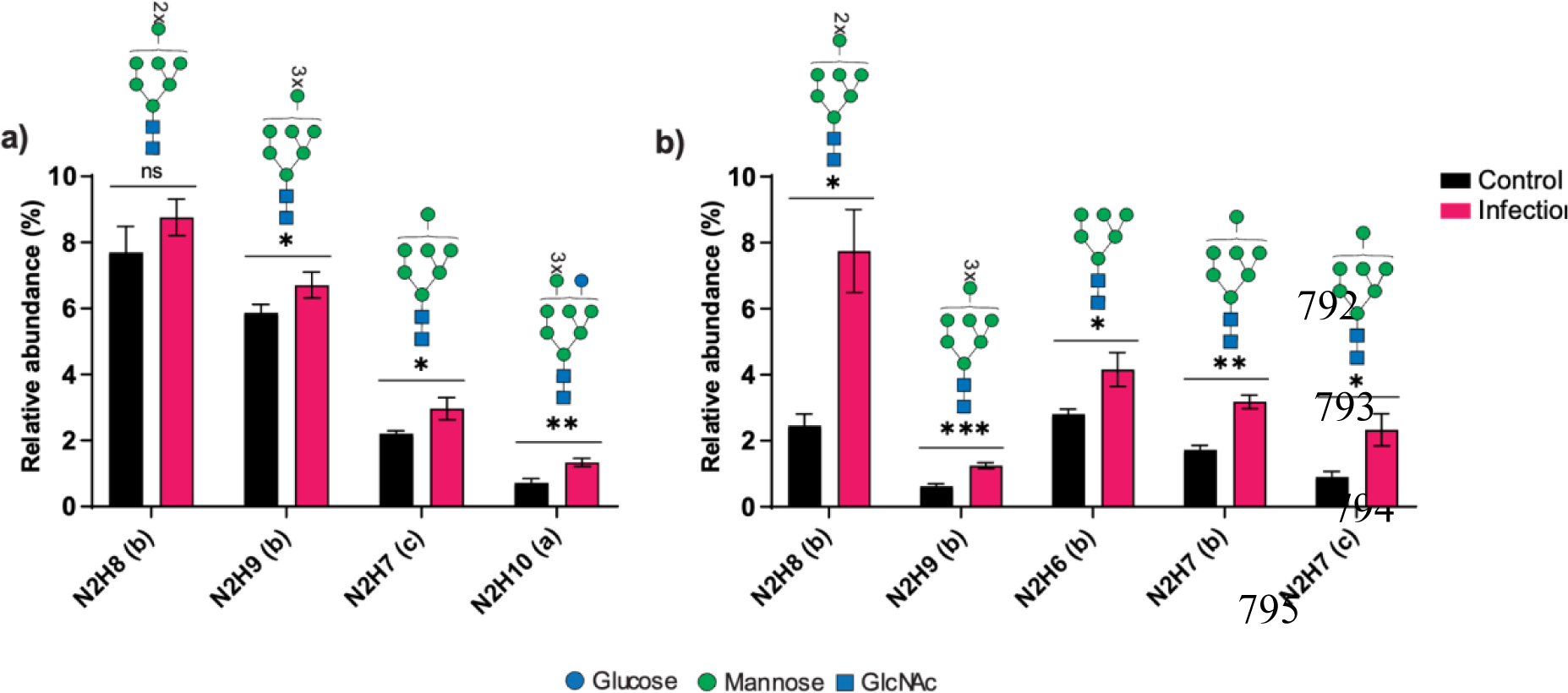
Alteration of oligomannose type *N*-glycan on mock-infected and infected cells at 48 h post-infection. Relative abundance of significantly altered oligomannose type glycans on a) cell extracted proteins, b) secreted proteins. Man: Mannose; Glc: Glucose. n=3, *P<0.05, **P<0.01, ns: not significant, unpaired Welch’s t-test.

**Fig S3:**
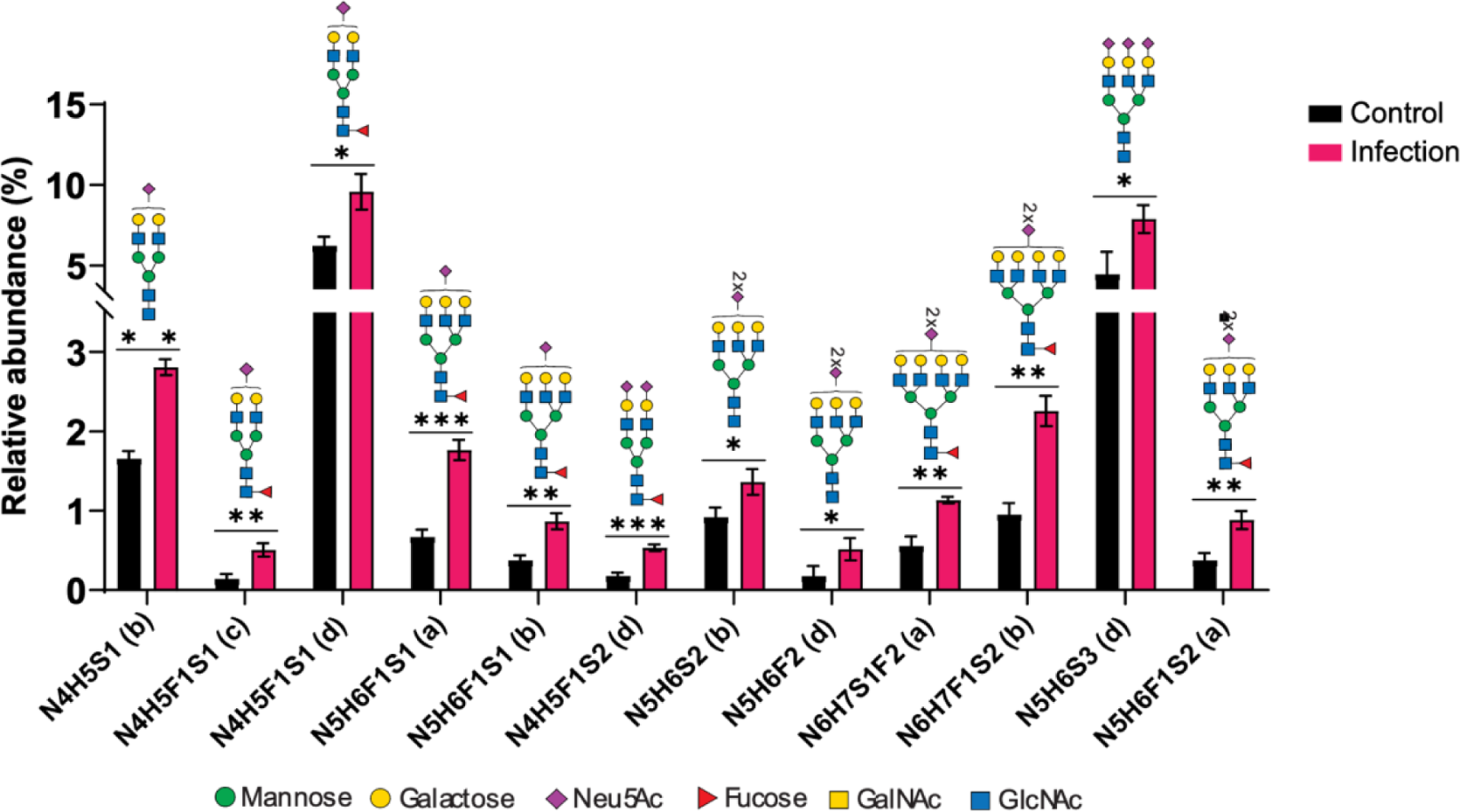
Significantly altered (increased expression) sialylated complex-type *N*-glycan species on cell extracted proteins at 24h post-infection. n=3, *P<0.05, **P<0.01, ***P<0.001, unpaired Welch’s t-test. N: *N*-acetylhexosamine; H: Hexose; F: Fucose; S: *N*-acetylneuraminic Acid.

**Fig S4:**
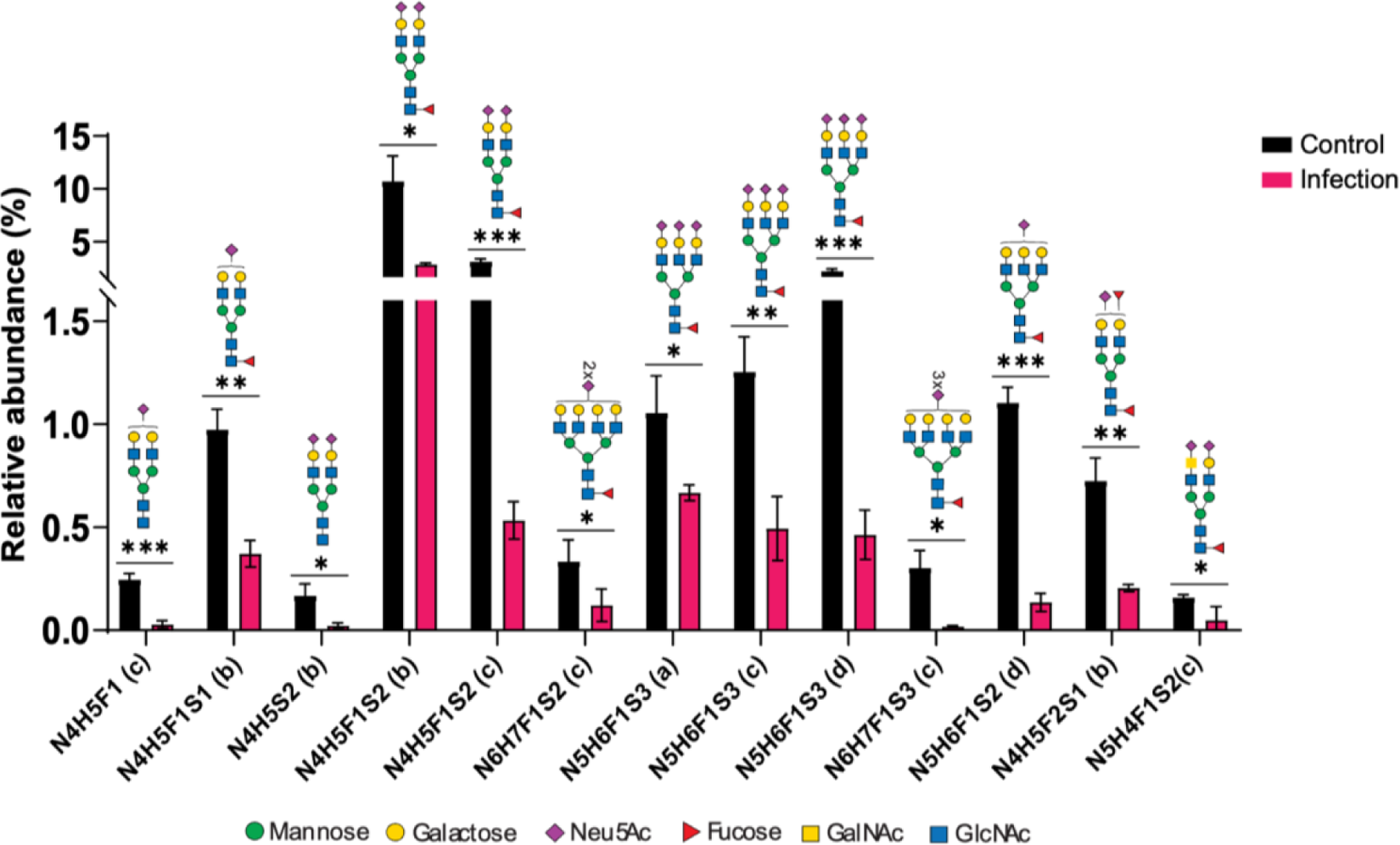
Significantly altered (decreased expression) sialylated complex-type *N*-glycan species on cell extracted proteins at 24h post-infection. n=3, *P<0.05, **P<0.01, ***P<0.001, unpaired Welch’s t-test. N: *N*-acetylhexosamine; H: Hexose; F: Fucose; S: *N*-acetylneuraminic Acid.

**Fig S5:**
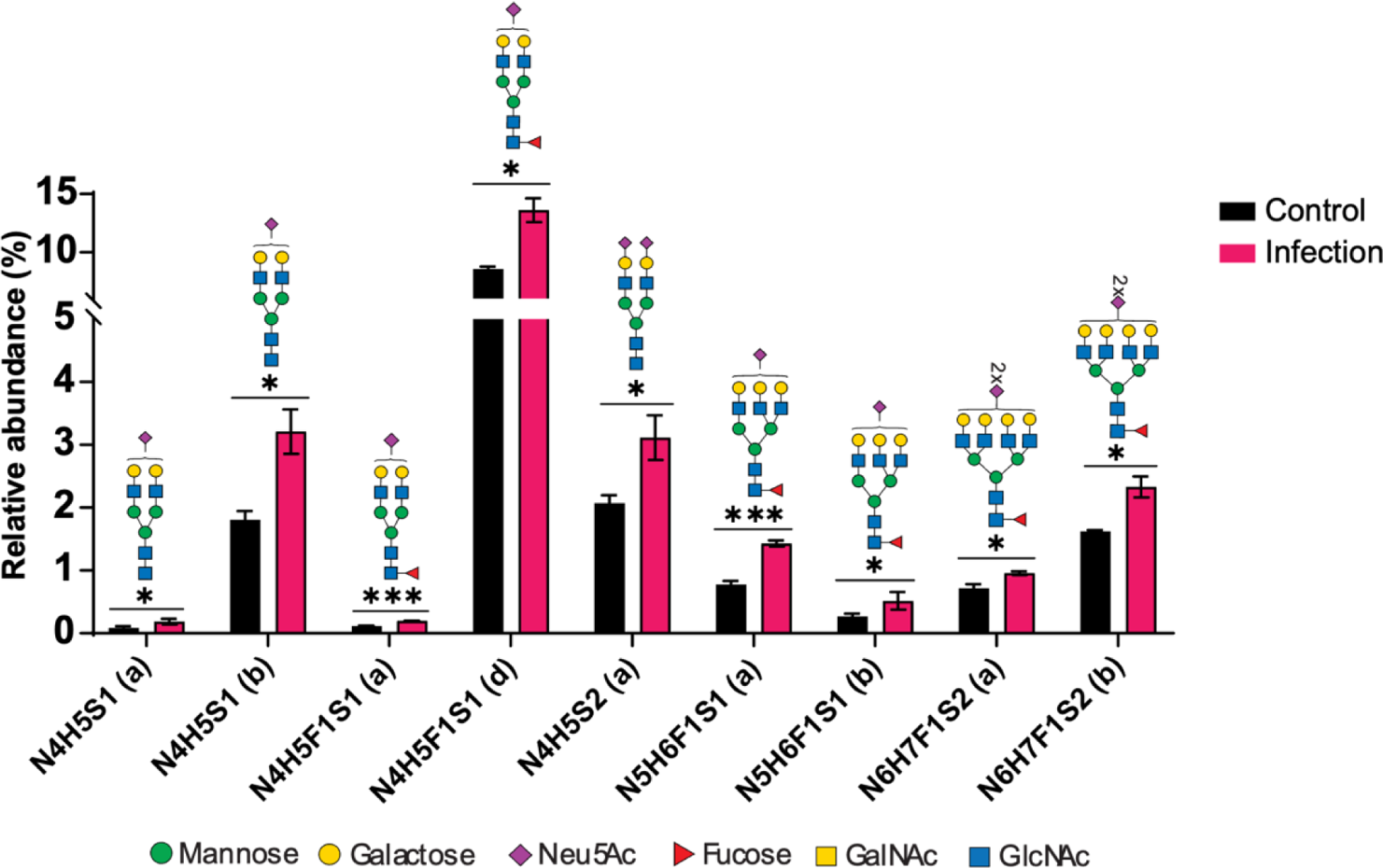
Significantly altered (increased expression) sialylated complex-type *N*-glycan species on secreted proteins at 24h post-infection. n=3, *P<0.05, **P<0.01, ***P<0.001, unpaired Welch’s t-test. N: *N*-acetylhexosamine; H: Hexose; F: Fucose; S: *N*-acetylneuraminic Acid.

**Fig S6:**
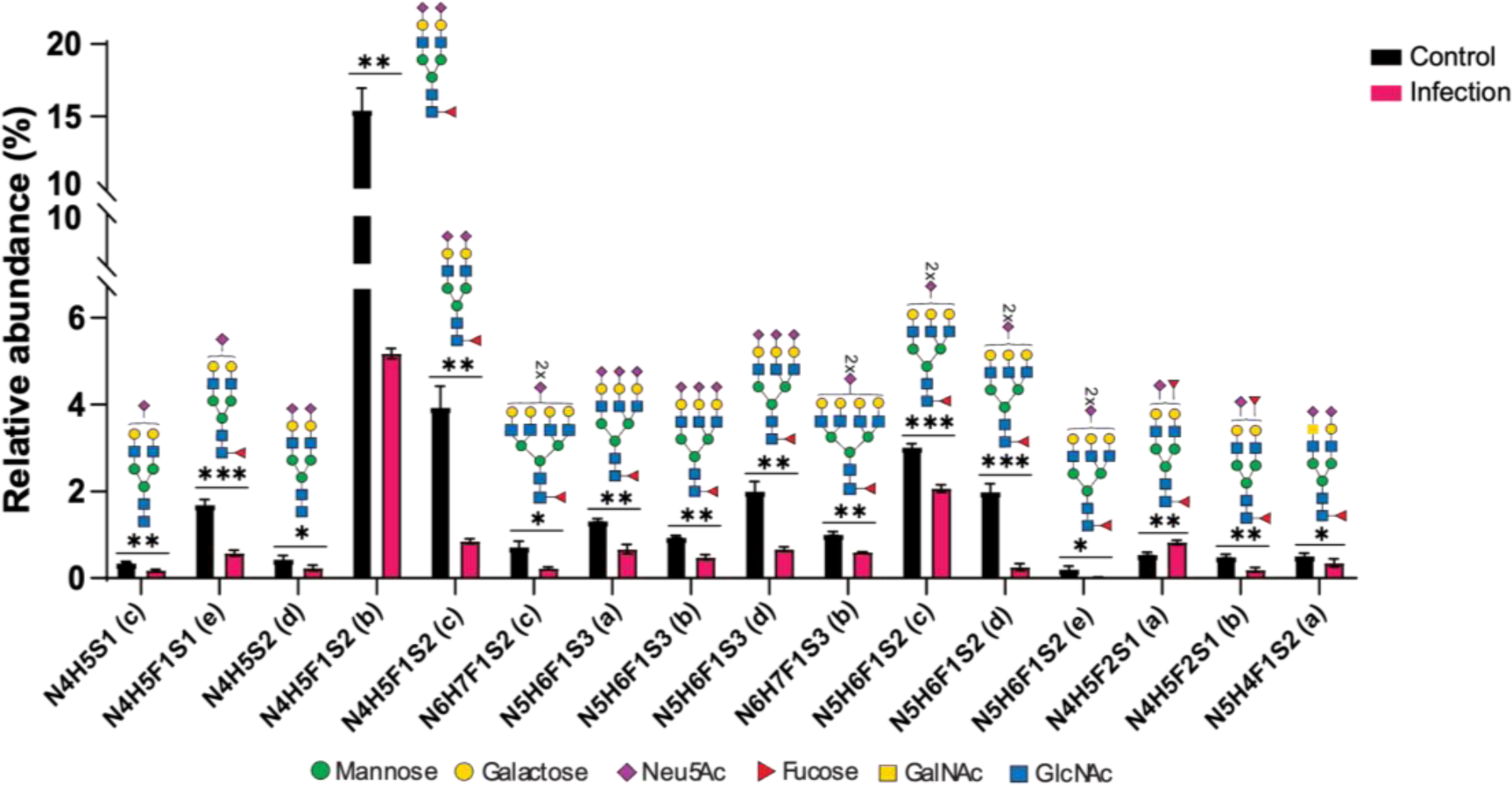
Significantly altered (decreased expression) sialylated complex-type *N*-glycan species on secreted proteins at 24h post-infection. n=3, *P<0.05, **P<0.01, ***P<0.001, unpaired Welch’s t-test. N: *N*-acetylhexosamine; H: Hexose; F: Fucose; S: *N*-acetylneuraminic Acid.

**Fig S7:**
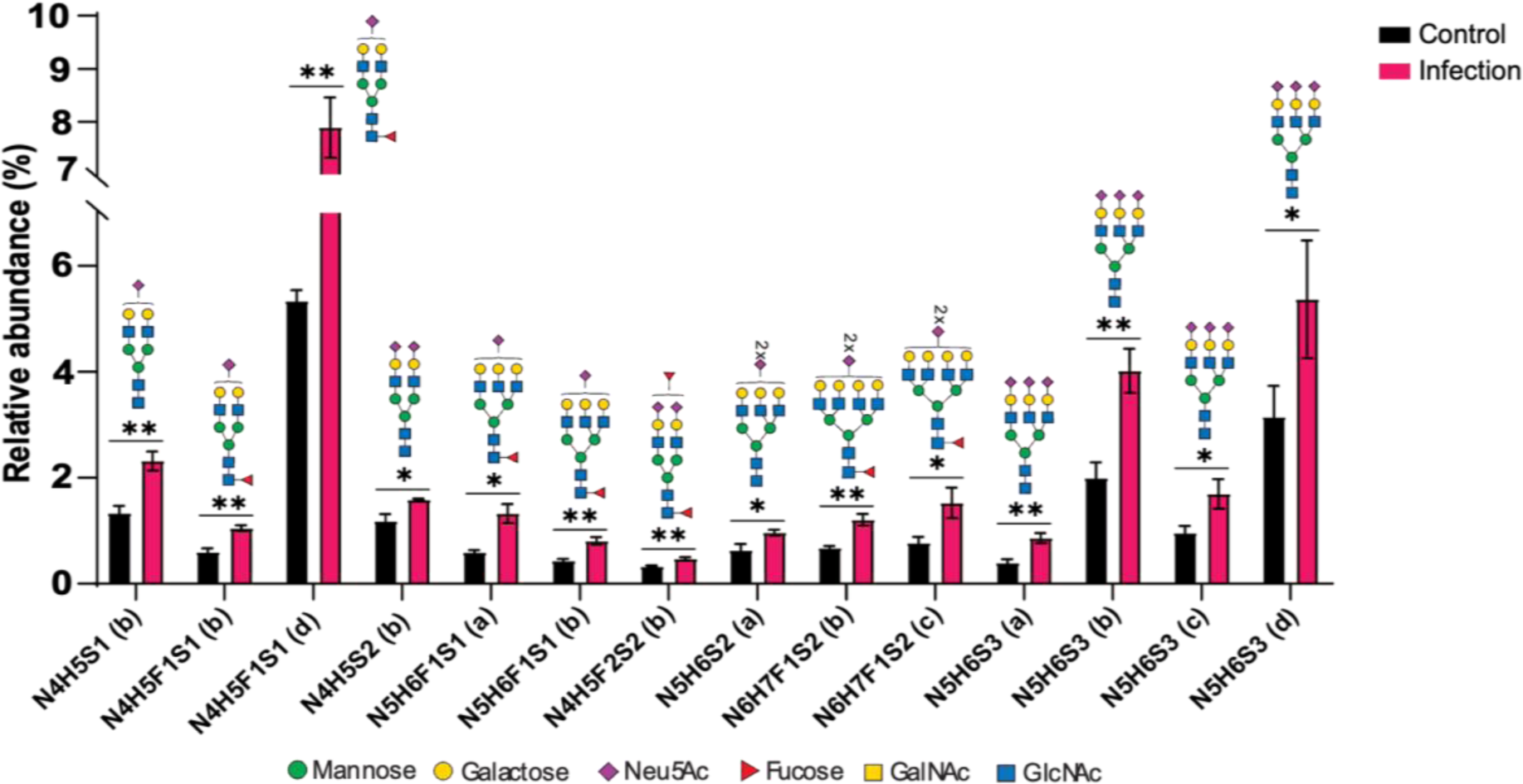
Significantly altered (increased expression) sialylated complex-type *N*-glycan species on cell extracted proteins at 48h post-infection. n=3, *P<0.05, **P<0.01, ***P<0.001, unpaired Welch’s t-test. N: *N*-Acetyl Hexosamine; H: Hexose; F: Fucose; S: *N*-acetylheuraminic Acid.

**Fig S8:**
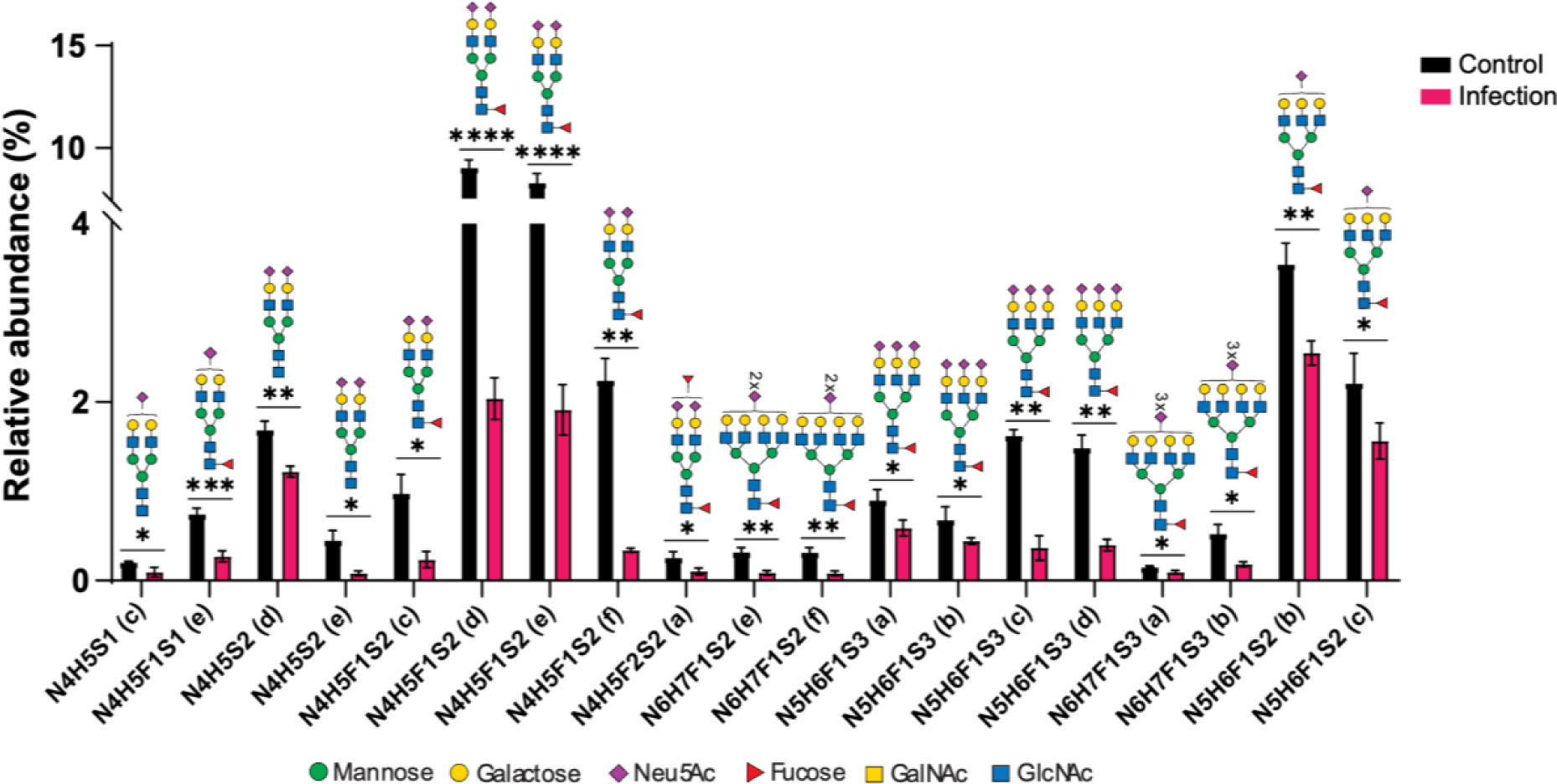
Significantly altered (decreased expression) sialylated complex-type *N*-glycan species on cell extracted proteins at 48h post-infection. n=3, *P<0.05, **P<0.01, ***P<0.001, ****P<0.0001, unpaired Welch’s t-test. N: *N*-acetylhexosamine; H: Hexose; F: Fucose; S: *N*-acetylneuraminic Acid.

**Fig S9:**
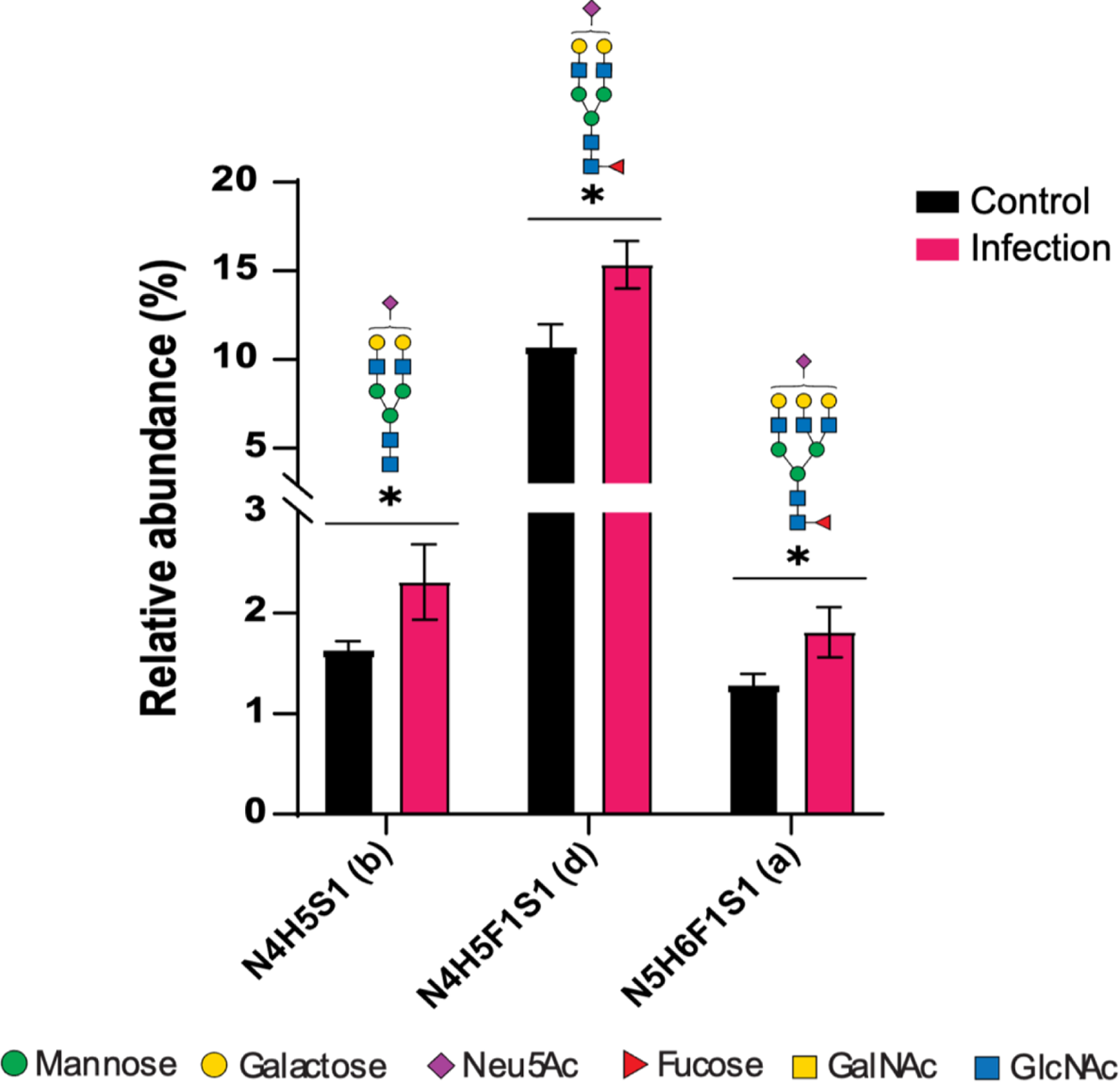
Significantly altered (increased expression) sialylated complex-type *N*-glycan species on secreted proteins at 48h post-infection. n=3, *P<0.05, **P<0.01, unpaired Welch’s t-test. N: *N*-acetylhexosamine; H: Hexose; F: Fucose; S: *N*-acetylneuraminic Acid.

**Fig S10:**
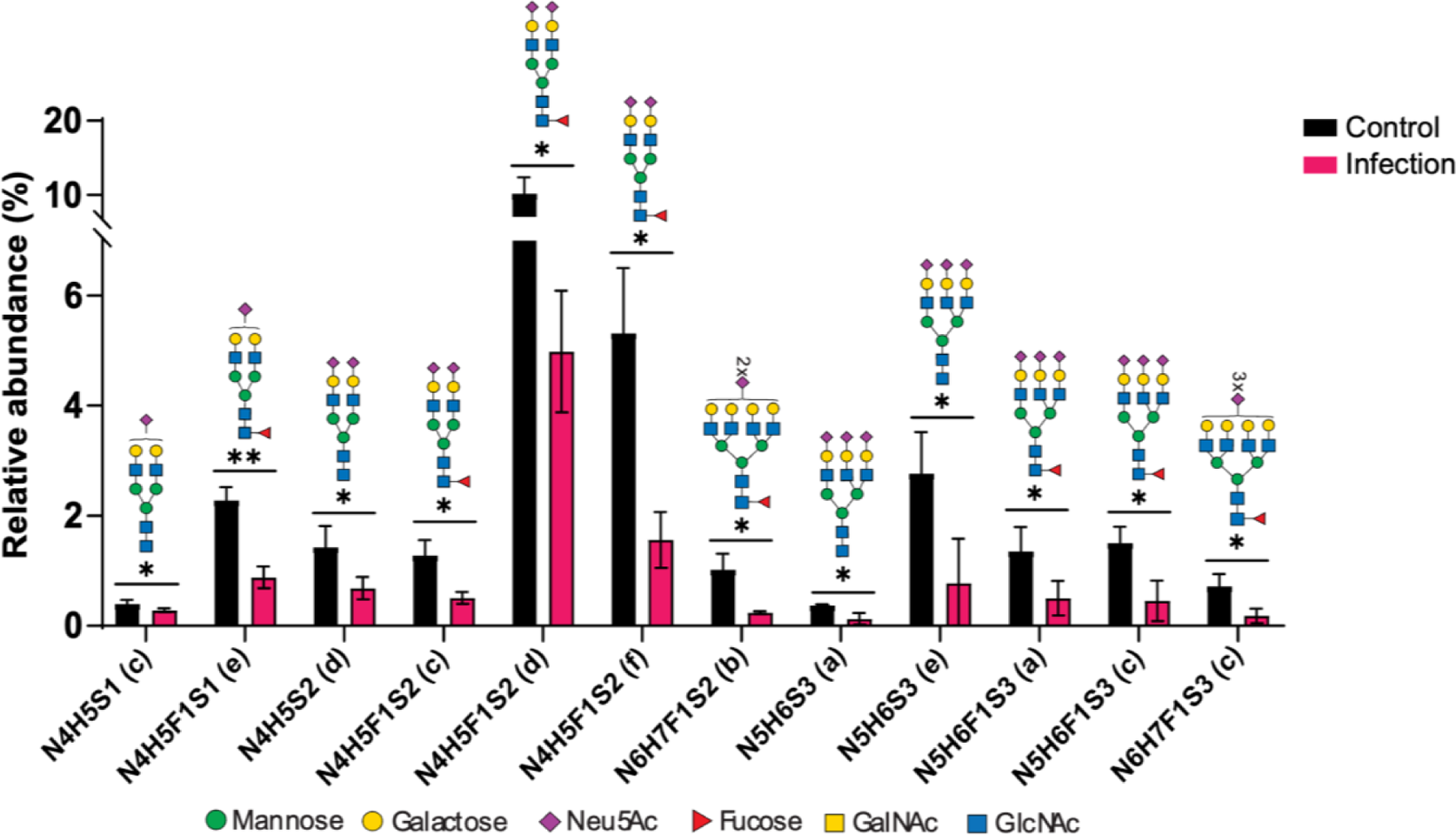
Significantly altered (decreased expression) sialylated complex-type *N*-glycan species on secreted proteins at 48h post-infection. n=3, *P<0.05, **P<0.01, unpaired Welch’s t-test. N: *N*-acetylhexosamine; H: Hexose; F: Fucose; S: *N*-acetylneuraminic Acid.

